# Phylogenomic Analysis of Protein-Coding Genes Resolves Complex Gall Wasp Relationships

**DOI:** 10.1101/2022.06.20.496719

**Authors:** Jack Hearn, Erik Gobbo, José Luis Nieves-Aldrey, Antoine Branca, James A. Nicholls, Georgios Koutsovoulos, Nicolas Lartillot, Graham N. Stone, Fredrik Ronquist

## Abstract

The phylogeny of gall wasps (Cynipidae) and their parasitic relatives has attracted considerable attention in recent years. The family is now widely recognized to fall into thirteen natural lineages, designated tribes, but the relationships among them have remained elusive. This has stymied any progress in understanding how cynipid gall inducers evolved from insect parasitoids, and what role inquilinism (development as a herbivore inside galls induced by other cynipids) might have played in this transition. A recent analysis of ultraconserved elements (UCEs) represents the first attempt at resolving these questions using phylogenomics. Here, we present the first analysis based on protein-coding sequences from genome and transcriptome assemblies. To address potential problems due to model misfit, we focus on models that accommodate site-specific amino-acid profiles and that are less sensitive than standard models to long-branch attraction. Our results show that the Cynipidae as previously circumscribed are not monophyletic. Specifically, the Paraulacini and a clade formed by Diplolepidini + Pediaspidini both fall outside a core clade (Cynipidae s. str.), which is more closely related to Figitidae. This result is robust to the exclusion of long-branch taxa that could potentially mislead the analysis, and it is consistent with the UCE analysis. Given this, we propose that the Cynipidae be divided into three families: the Paraulacidae, Diplolepididae and Cynipidae (s. str.). Our results suggest that the Eschatocerini are the sister group of the remaining Cynipidae (s. str.). Within the latter, our results are consistent with the UCE analysis but place two additional tribes: (1) the Aylacini (s. str.), more closely related to the oak gall wasps (Cynipini) and some of their inquilines (Ceroptresini) than to other herb gallers (Aulacideini and Phanacidini); and (2) the Qwaqwaiini, likely the sister group to Synergini (s. str.) + Rhoophilini. Several alternative scenarios for the evolution of cynipid life histories are compatible with the relationships suggested by our analysis, but all are complex and require multiple shifts between parasitoids, inquilines and gall inducers. Linking the different types of life-history transitions to specific genomic signatures may be one of the best ways of differentiating among these alternative scenarios. Our study represents the first step towards enabling such analyses.

## Introduction

Gall wasps (Hymenoptera: Cynipidae) induce the development of highly modified plant tissues, termed galls, in which their immature stages develop (Melika & Abrahamson 2002; Stone et al. 2002). The cynipid larva is enclosed inside a gall chamber lined with specialised nutritive cells formed by the plant in response to signals released by the gall wasp egg and larva (Stone & Schönrogge 2003; Harper et al. 2004; Hearn et al. 2019). While all cynipids appear able to induce the development of such nutritive tissues, several lineages – termed inquilines – can only induce nutritive tissue development within galls initiated by other species (Sanver & Hawkins 2000). The inquilines can thus be seen as cynipids that induce a ’gall within a gall’. The presence of inquilines can negatively affect the fitness of the primary gall-inducer, in many cases killing them (László & Tóthmérész 2006). In *Periclistus* inquilines, it has been reported that the ovipositing female kills the larva of the gall inducer by stabbing it with her ovipositor, potentially injecting harmful substances in the process (Shorthouse 1973).

Several hypotheses on the mechanism of cynipid gall induction have been advanced, partly inspired by knowledge of other gall-inducing organisms: secretion of auxins (Tooker & Helms 2014), injection of virus-like particles (Cornell 1983; Cambier et al. 2019), manipulation of plant NOD factors (Hearn et al. 2019), or involvement of bacterial or fungal symbionts (Hearn et al. 2019). However, in contrast to some other gall induction systems (Harris & Pitzschke 2020), there is no conclusive evidence for any of these hypotheses in cynipids.

Our understanding of the evolutionary origin of cynipid gall inducers and inquilines is equally poor. The Cynipidae are deeply nested within the insect-parasitic Apocrita (Ronquist 1995, 1999; Heraty et al. 2011; Sharkey et al. 2011; Klopfstein et al. 2013; Peters et al. 2017), and all other members of the superfamily Cynipoidea are insect parasitoids, so it has long been clear that the phytophagous gall inducers and inquilines must have evolved from insect-parasitic ancestors. It has generally been assumed that the phytophagous forms constitute a monophyletic lineage in the Cynipoidea, the family Cynipidae, although it has been surprisingly difficult to find morphological characters supporting their monophyly (Liljeblad & Ronquist 1998; Ronquist 1999; Ronquist et al. 2015).

Since Ashmead (1903), the Cynipidae have commonly been divided into six tribes: the Cynipini, Diplolepidini (or Rhoditini), Pediaspidini, Eschatocerini, Aylacini and Synergini. The Cynipini comprise the oak gall inducers, one of the largest radiations of insect gall inducers with more than 1,000 described species, most of which are associated with oaks (*Quercus*). The Diploplepidini consist of the gall inducers on roses (*Rosa*), among them the well-known bedeguar gall wasp, *Diplolepis rosae*. The Pediaspidini and Eschatocerini are two small tribes, originally including a single genus each: *Pediaspis*, a European genus inducing galls on maples (*Acer*), and *Eschatocerus*, a South American gall inducer on *Vachellia* (commonly known as thorn trees or acacias) and other woody members of Fabaceae. The inquilines are grouped in this system into the Synergini, and the remaining gall inducers, mostly associated with herbaceous host plants, in the Aylacini.

Early analyses of cynipid relationships based on morphological data suggested that the Aylacini form a paraphyletic assemblage of early-diverging cynipid lineages (Ronquist 1994; Liljeblad & Ronquist 1998), consistent with ideas presented over a century ago by the famous cynipidologist and later sexologist Alfred Kinsey (Kinsey 1920). They also suggested that the oak gall wasps (Cynipini) and the inquilines (Synergini) form natural monophyletic groups. The former appeared to be related to the Diplolepidini, Eschatocerini and Pediaspidini, all inducing galls on trees or bushes belonging to the rosid clade of angiosperms (“woody rosids”). An important result was that the genus *Himalocynips*, a cynipid from Nepal with unknown biology and originally placed in a separate subfamily (Yoshimoto 1970), was grouped with *Pediaspis* (Liljeblad and Ronquist 1998). These early analyses also indicated that the inquilines (Synergini) originated from gall inducers related to the Aylacini genera *Diastrophus* and *Xestophanes* (Ronquist 1994; Liljeblad & Ronquist 1998).

Subsequent analyses of molecular data and combined molecular, morphological and life-history data (Nylander et al. 2004; Ronquist et al 2015) have confirmed some of these results and rejected others. Among the inquilines, only *Periclistus* (inquilines in *Diplolepis* galls on *Rosa*) and *Synophromorpha* (inquilines in *Diastrophus* galls on *Rubus*) appear to be closely related to *Diastrophus* and *Xestophanes*. Together, they form a strongly supported lineage of gall inducers and inquilines associated with herbaceous and woody hosts in the Rosaceae, now recognized as the tribe Diastrophini (Ronquist et al 2015; Table 1). The remaining Aylacini appear to fall into three distinct lineages that are now recognized as separate tribes (Ronquist et al 2015): (1) the Aylacini (s. str.), gall inducers associated with poppies (*Papaver*); (2) the Aulacideini, gall inducers mostly associated with Asteraceae and Lamiaceae, but also with a few other families, including Papaveraceae; and (3) the Phanacidini, gall inducers mainly associated with Asteraceae and Lamiaceae, and often inducing stem galls. The remaining inquilines appear to fall into two distinct monophyletic lineages (Ronquist et al. 2015): (1) the Ceroptresini, including the single genus *Ceroptres* associated with Cynipini oak galls; (2) and a lineage consisting of Synergini (s. str.), comprising the remaining inquilines of Cynipini oak galls, and *Rhoophilus*, an inquiline in lepidopteran galls on species of *Searsia* (Anacardiaceae) (van Noort et al. 2007). Several analyses have supported a sister-group relationship between *Rhoophilus* and the remaining Synergini (s. str.) (Liljeblad & Ronquist 1998; Ronquist et al. 2015; Ide et al. 2018) and recently it was proposed to recognize a separate tribe for *Rhoophilus*, the Rhoophilini (Lobato-Vila et al. 2022).

**Table 1.** Life history of the 13 tribes of Cynipidae recognized currently (Ronquist et al. 2015; Lobato-Vila et al. 2022).

| Tribe | Life history | Host |
| --- | --- | --- |
| Aulacideini | Gall inducers | Herbs in the families Asteraceae, Lamiaceae, and others |
| Aylacini (s. str.) | Gall inducers | Poppies (Papaveraceae) |
| Ceroptresini | Inquilines | Galls of Cynipini on oaks |
| Cynipini | Gall inducers | Oaks, occasionally related trees (Fagaceae) |
| Diastrophini | Gall inducers and inquilines | Gall inducers on Rosaceae (*Rubus* and *Potentilla*), and inquilines in cynipid galls on Rosaceae (*Rubus* and *Rosa*) |
| Diplolepidini | Gall inducers | Roses (Rosaceae) |
| Eschatocerini | Gall inducers | *Vachellia*, *Prosopis* (Fabaceae) |
| Paraulacini | Parasitoids (see text) | *Aditrochus* (Pteromalidae) gall inducers on *Nothofagus* (Nothofagaceae, Fagales) |
| Pediaspidini | Gall inducers | Maples (Sapindaceae) |
| Phanacidini | Gall inducers | Herbs, mostly in the families Asteraceae and Lamiaceae |
| Qwaqwaiini | Gall inducers | *Scolopia* (Salicaceae) |
| Rhoophilini | Inquilines | Lepidoptera galls on *Searsia* (Anacardiaceae) |
| Synergini (s. str.) | Inquilines; a few gall inducers | Galls of Cynipini; a few are true gall inducers on oaks (Fagaceae) |

Among the species associated with woody rosids, the molecular analyses have clearly supported the monophyly of the oak gallers (Cynipini) and rose gallers (Diplolepidini; Table 1). In recent years, two additional lineages associated with woody rosids have been added, the Qwaqwaiini and Paraulacini (Table 1). The tribe Qwaqwaiini is based on a newly discovered gall inducer on *Scolopia* (Salicaceae) in South Africa (Liljeblad et al. 2011), while the Paraulacini constitute a re-discovered lineage of cynipids associated with galls on southern beeches (*Nothofagus*; Nothofagaceae) (Nieves-Aldrey et al. 2009).

In conclusion, this results in the current classification of the Cynipidae into 13 tribes (Table 1) (Ronquist et al. 2015; Lobato-Vila et al. 2022). The analyses described above, based only on a few molecular markers, have been unable to resolve the relationships between these 13 lineages, with three notable exceptions. First, there has been fairly strong evidence for a sister-group relationship between the Diplolepidini and Pediaspidini, suggesting that at least these two lineages of well-known woody-rosid gallers are related (Ronquist et al. 2015). Second, there has been support for a sister-group relationship between the two major lineages of herb gallers, the Aulacideini and Phanacidini (Nylander et al. 2004; Ronquist et al. 2015). Third, there is strong support for the sister-group relationship between the Rhoophilini and Synergini (s. str.), as mentioned above. Many of the tribes, however, appear to represent isolated lineages, with no close relatives among the other tribes (Nylander et al. 2004; Ronquist et al. 2015).

Regardless of the relationships among the major lineages, these findings have made it clear that the evolutionary origin of cynipid gall inducers and inquilines is more complicated than originally thought. In the tribe Diastrophini, for instance, there are two genera of inquilines and two genera of gall inducers, and the current data indicate that there must have been at least two transitions between these life-history strategies within the tribe (Ronquist et al 2015). Recent work has also shown that the tribe Synergini (s. str.), previously considered to consist entirely of inquilines, includes at least one deeply nested lineage of true gall inducers—*Synergus itoensis* and related species—inducing galls inside acorns (Abe, Ide, and Wachi 2011; Ide et al. 2018; Gobbo et al. 2020). There are also observations suggesting that facultative intraspecific inquilinism may occur in *Diastrophus* (Diastrophini) (Pujade i Villar 1984), and it has been suggested that the remarkable parallelisms between the Aulacideini and Phanacidini in the evolution of host plant preferences could be due to facultative or obligate inquilinism among some cynipid herb gallers (Ronquist & Liljeblad 2001; Nieves-Aldrey, Gómez & Hernández Nieves 2004).

The recent discovery of gall-inducing Synergini (s. str.) illustrates how difficult it is to correctly deduce the life history of insects reared from galls, and previous assumptions about the life history of different cynipid lineages should be analyzed critically. The Paraulacini are a case in point. Recent studies have revealed that they are associated with *Nothofagus* galls that are presumably induced by the chalcidoid genus *Aditrochus*, currently placed in the Pteromalidae, but it has remained unclear whether the Paraulacini are phytophagous inquilines or parasitoids of some other gall inhabitant (Nieves-Aldrey et al. 2009; Ronquist et al. 2015). The relatively close relationship between southern beeches (*Nothofagus*) and oaks, both belonging to the Fagales, would suggest that the Cynipini and Paraulacini might also be related, and both phytophagous. However, recent genetic analyses have provided a case where the genome of *Cecinothofagus ibarrai* (Paraulacini) was retrieved from a larva of *Aditrochus coihuensis*, together with the *Aditrochus coihuensis* genome (Rasplus, Nieves-Aldrey & Cruaud 2022). This suggests that the Paraulacini are not only parasitoids, they are also likely koinobiont endoparasitoids in early larval instars, like all other insect-parasitic cynipoids (see below).

The life-history of the Eschatocerini is another interesting case. Members of this tribe have been reared from galls on *Prosopis* and *Vachellia* (formerly *Acacia*) collected in Argentina and Chile, and they have been assumed to be gall inducers (Nieves-Aldrey & San Blas 2015; Aranda-Rickert et al. 2017). However, like the *Nothofagus* galls, these galls also produce a number of other insects that could potentially be gall inducers. These include *Allorhogas prosopidis* (Braconidae), a genus of phytophagous braconids that may be inquilines or gall inducers (Samacá-Sáenz, Egan & Zaldívar-Riverón 2020), and the chalcidoid *Tanaostigmus coeruleus* (Chalcidoidea, Tanaostigmatidae), which belongs to a genus that is known to include phytophagous species (either inquilines or gall inducers). The galls are also inhabited by members of several genera of eurytomids, namely *Proseurytoma, Sycophila* and *Eurytoma*. Other members of these genera include true gall inducers, such as *Proseurytoma gallarum*, a gall inducer on *Geofreoa decorticans* (another Fabaceae sharing habitats with *Prosopis* and *Acacia*). Preliminary data available to one of us (JLNA) suggest that *Allorhogas* and *Tanaostigmodes* are both inquilines in *Eschatocerus* galls, which is at least consistent with *Eschatocerus* being the true gall inducer. Here, we will assume that the Eschatocerini are gall inducers, but additional evidence supporting this conclusion would be highly desirable. Among the remaining cynipid tribes, we still lack detailed studies of the life history for the Ceroptresini (apparently inquilines; but see Blair 1949) and Qwaqwaiini (apparently true gall inducers; Liljeblad et al. 2011).

Except for the Cynipidae, the superfamily Cynipoidea comprises the families Austrocynipidae, Ibaliidae, Liopteridae, and Figitidae (Ronquist 1995; 1999). The life history of several species of ibaliids and figitids is well-studied (Ronquist 1999, and references cited therein). They are all koinobiont endoparasitoids in early larval instars. Towards the end of their development, they emerge and consume the remains of the moribund host as ectoparasitoids. The most diverse lineage is the Figitidae, which has appeared as the sister group of the Cynipidae in most previous analyses (Ronquist 1995; Ronquist 1999; Buffington, Nylander & Heraty 2007; Buffington et al. 2012; Ronquist et al. 2015).

The origin of the Cynipidae appears to be linked to that of several lineages of gall-associated Figitidae, which appear to form early-diverging lineages in the family (Ronquist 1995; Ronquist 1999; Buffington, Nylander & Heraty 2007, Buffington et al. 2012; Ronquist et al 2015). These gall-associated figitids include the Parnipinae (Ronquist and Nieves-Aldrey 2001), Plectocynipinae (Ros-Farré & Pujade-Villar 2007; Buffington & Nieves-Aldrey 2011), Thrasorinae (Buffington 2008; Paretas-Martínez et al 2011), Mikeiinae (Paretas-Martínez et al 2011) and Euceroptresinae (Buffington & Liljeblad 2008; note that the subfamily name should be Euceroptresinae and not Euceroptrinae). There is fairly strong evidence that the Parnipinae are koinobiont early-internal-late-external parasitoids of cynipid gall inducers in the genera *Barbotinia* and *Iraella* (Ronquist et al. 2018). The life-history of the other lineages remains unclear, although they are generally assumed to be parasitoids of other inhabitants in the cynipid and chalcidoid galls from which they have been reared.

A recent paper represents the first attempt at resolving phylogenetic relationships in the Cynipoidea using phylogenomic data (Blaimer et al. 2020). Specifically, this analysis used an approach known as ultra-conserved elements (UCEs) to obtain genomic data from a wide range of cynipoid exemplars, representing all families except the Austrocynipidae and spanning a significant amount of the known diversity within each family. Several surprising results emerged from this UCE analysis. First, the Liopteridae and Ibaliidae were placed within the Figitidae, among the early-diverging gall-associated lineages. Second, the Paraulacini and the Diplolepidini + Pediaspidini were placed outside the clade formed by the Figitidae and the remaining Cynipidae (the Cynipidae s. str.) - a relationship first hinted at in Hymenoptera-wide analyses (Peters et al. 2017). Finally, the analysis suggested that the Eschatocerini may be the sister group of the Figitidae, although the evidence for this was weak and alternative placements appeared under some analysis settings.

The analysis we present here is the first phylogenomic analysis based on genome and transcriptome assemblies, and it allows a largely independent test of the results from the UCE analysis. In contrast to the UCE analysis, our taxon sampling is focused on cynipids. It lacks ibaliids and liopterids, and is relatively sparse with respect to figitids. However, it includes representatives of all cynipid tribes except the recently recognized Rhoophilini. Importantly, it includes the Qwaqwaiini and Aylacini (s. str.), both of which were missing from the UCE analysis. The UCE study claims to include one representative of the Aylacini (s. str.), *Aylax salviae*. However, this species has long been placed in the genus *Neaylax* (Nieves-Aldrey 1994, 2001), which is deeply nested inside the Aulacideini (Ronquist et al. 2015). Specifically, *Neaylax salviae* belongs to a clade of Aulacideini gallers of Lamiaceae related to the genus *Antistrophus* (Ronquist et al. 2015), and this is entirely consistent with the placement of *“Aylax” salviae* in the UCE analysis (Blaimer et al. 2020). Another key taxon represented in our analysis but not in the UCE analysis is the single species in the recently described genus *Protobalandricus, P. spectabilis,* which represents a divergent sister group to all other sampled Cynipini (Nicholls, Stone & Melika 2018).

Importantly, by focusing on data from protein-coding genes, we can use sophisticated substitution models that accommodate variation in amino-acid profiles across sites. These models are known to resolve some issues with long-branch attraction that can affect analyses under standard models, such as those used in the UCE analysis (Kapli and Telford 2020). The most surprising UCE results do involve the placement of long, isolated lineages—the Paraulacini, Eschatocerini and Diplolepidini + Pediaspidini—so there is reason to suspect that such phenomena may be at play. Based on the results of our analysis, which largely confirm and complement the UCE results, we propose a new family-level classification of the Cynipidae. We also discuss the implications with respect to the evolutionary origin of cynipoid gall inducers and inquilines.

## Material and methods

### Taxon sampling

Species were chosen to represent all of the currently recognized tribes of cynipid gall wasps except Rhoophilini, and we tried to cover as much of the phylogenetic diversity within each lineage as possible (Table 2). Our Cynipini selection included *Protobalandricus spectabilis*, inducing galls on *Quercus* section *Protobalanus* oaks in California. The other species came from diverse Cynipini genera: *Andricus*, *Belonocnema*, *Biorhiza*, *Druon* and *Neuroterus*. In the Aulacideini, we included two gallers of Asteraceae (*Isocolus centaureae* and *Aulacidea tavakolii*), one galler of Lamiaceae (*Hedickiana levantina*) and one galler of Papaveraceae (*“Aylax” hypecoi*), thus covering much of the diversity in host-plant preferences in the group. The last species, *“Aylax” hypecoi*, is known to belong to the Aulacideini (Ronquist et al. 2015) even though its current generic placement suggests it is an Aylacini (s. str). Its relationships are such that it will most likely have to be placed in a new genus (Nieves-Aldrey in press); here, we will consistently use quotes around the genus name to denote that it is known not to belong to *Aylax*. Our Diastrophini selection included one inquiline (*Periclistus*) and one gall inducer (*Diastrophus*), covering both of the major life history strategies in the tribe. Our Synergini (s. str.) selection was unfortunately restricted to the most species-rich genus, *Synergus*, but it included both a gall inducer (*S. itoensis*) and three inquilines (*S. gifuensis*, *S. japonicus* and *S. umbraculus*). The remaining eight tribes were represented by single species; most of these tribes include few species and have uniform life histories (Table 1). The selection of exemplars for this study was completed before the appearance of the recent UCE study (Blaimer et al. 2020), but it does cover all major cynipid lineages detected in that analysis.

**Table 2.**
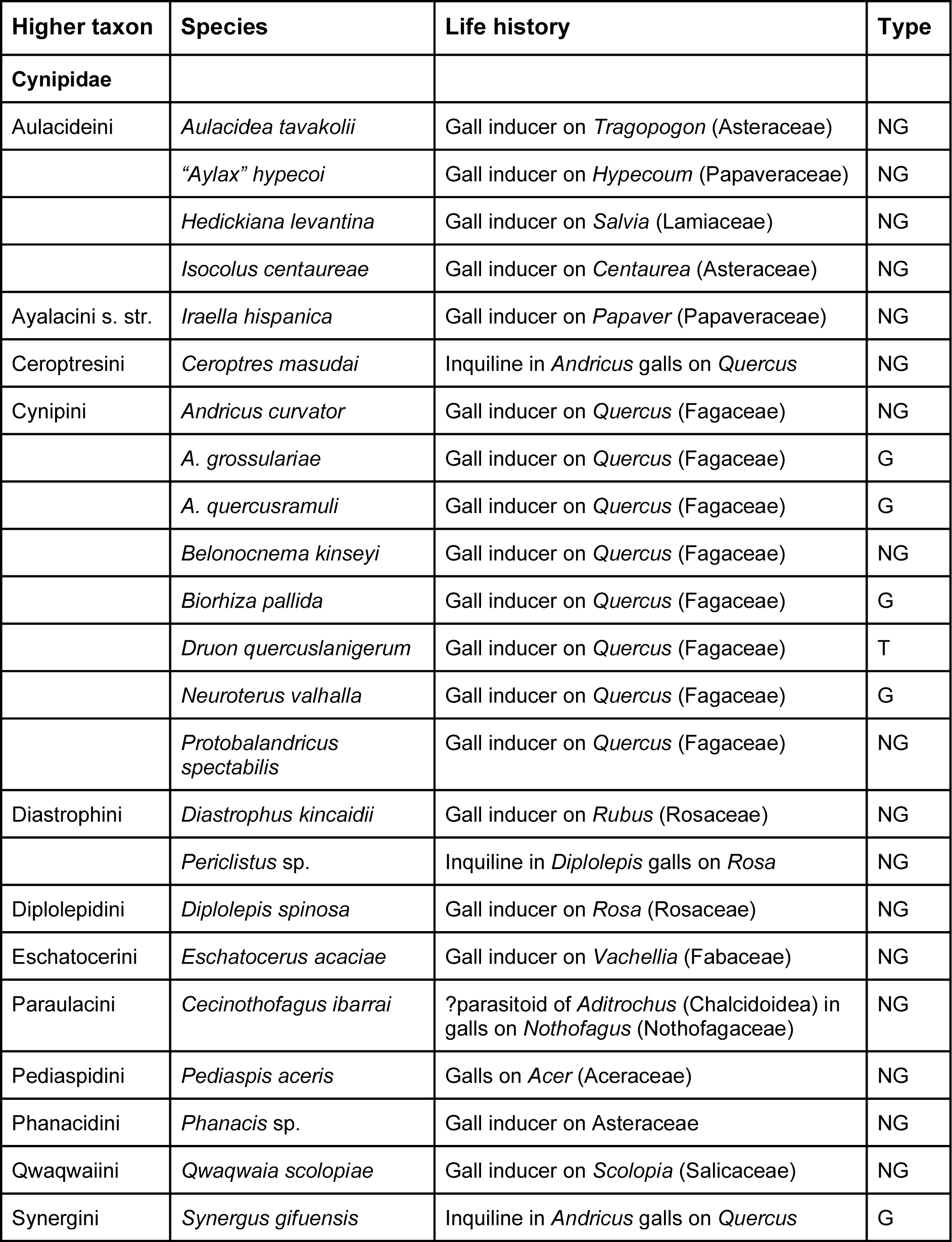
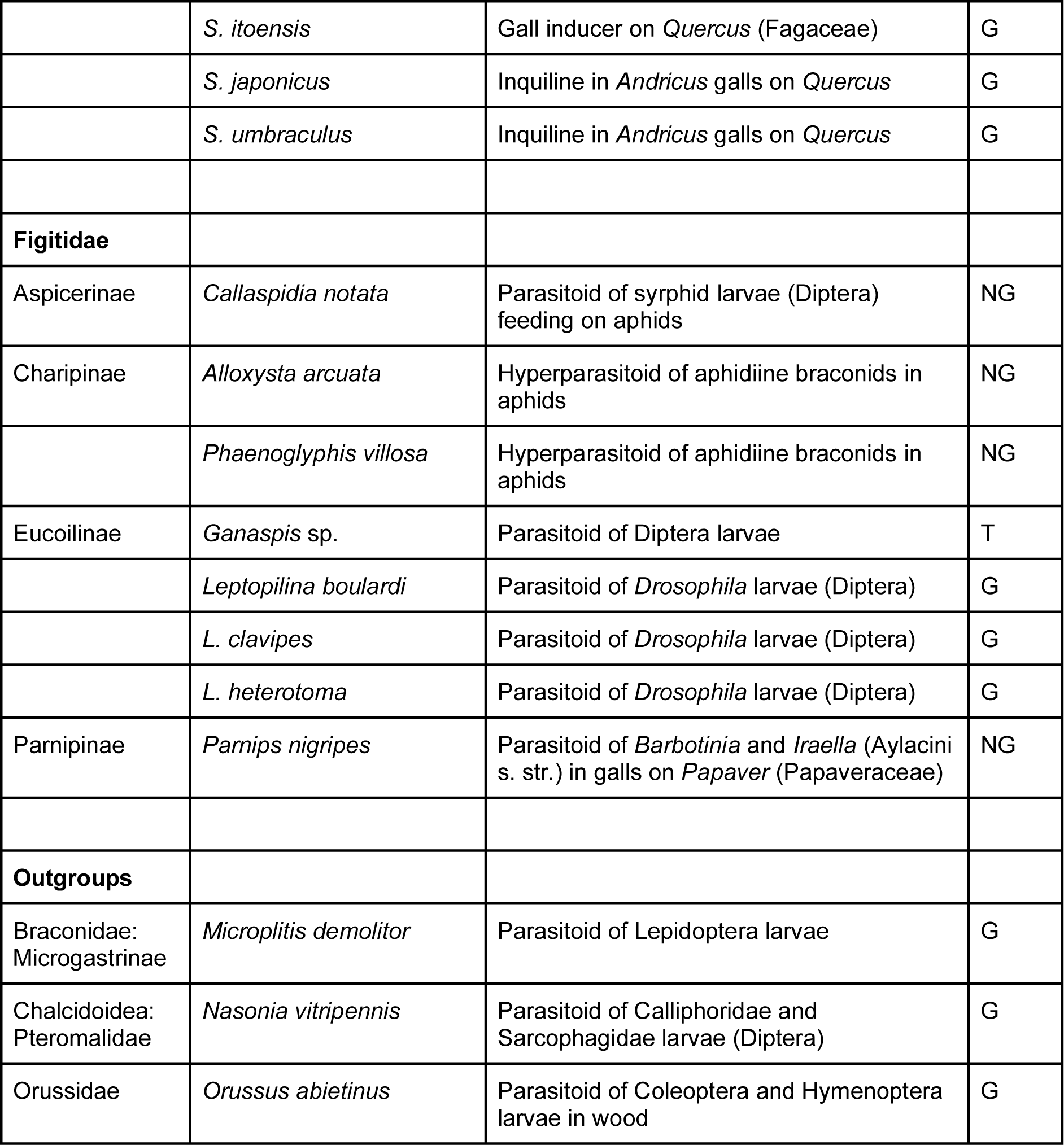
Overview of the species included in this study. For full data on the genome and transcriptome assemblies, see Supplementary Material, Table S1. Abbreviations for data type: G = Previously published genome assembly; NG = New genome assembly reported here, T = Previously published transcriptome. Note that the *Neuroterus valhalla* reference was recorded as *Callirhytis* sp. in the NCBI assembly, and *Belonocnema kinseyi* was recorded under the previous name *B. treatae* (Zhang et al. 2021).

Our sampling covers the entire diversity of Figitidae (Table 2). Importantly, our selection includes *Parnips nigripes* (Parnipinae), the only gall-associated figitid whose life history is known in some detail (Ronquist et al. 2018). The Parnipinae have appeared in previous analyses as the sister group of the remaining Figitidae, or even as the sister group of the Cynipidae (Ronquist 1999; Buffington, Nylander & Heraty 2007; Ronquist et al. 2015; Blaimer et al. 2020). We also included three more distant outgroups: *Orussus abietinus* (Orussidae), *Nasonia vitripennis* (Chalcidoidea: Pteromalidae) and *Microplitis demolitor* (Braconidae: Microgastrinae) (Table 2). Of those, *O. abietinus* is the most distant (Heraty et al. 2011; Sharkey et al. 2011; Klopfstein et al. 2013; Peters et al. 2017), and was used for rooting the trees generated in our analyses.

### Genome and transcriptome data

Two publicly available transcriptomes were included for the oak gall wasp *Biorhiza pallida* (Hearn et al. 2019), and the figitid *Ganaspis species 1* (Mortimer et al. 2013). Genome assemblies for the oak gall wasps *Andricus grossulariae*, *Belonocnema kinseyi*, *Druon quercuslanigerum* and *Neuroterus valhalla* (Brandão-Dias et al. 2022), four species of *Synergus* (Gobbo et al. 2020), three species of the figitid genus *Leptopilina* and three outgroups (*Nasonia*, *Microplitis* and *Orussus*) were downloaded from NCBI. The remaining data consisted of genomes assembled de novo for the current study (Table 2). References to all genome and transcriptome assemblies are provided in the Supplementary Material (Table S1).

### De novo genome assemblies

Two protocols were followed (Supplementary Material, Table S1). For the *Andricus curvator* and *A. quercusramuli* assemblies, DNA was extracted from single adults using the Thermo Scientific KingFisher Cell and Tissue DNA Kit and the KingFisher Duo magnetic particle processor. Genomes were sequenced by the Swedish National Genomic Infrastructure from ChromiumX libraries (Zheng et al. 2016) on a NovaSeq6000 (NovaSeq Control Software 1.6.0/RTA v3.4.4) with a 2x151 setup using ’NovaSeqXp’ workflow in ’S4’ mode flow cell. The Bcl to FastQ conversion was performed using bcl2fastq_v2.20.0.422 from the CASAVA software suite. Filtering and assembly were done by running 10X Genomics’ Supernova version 2.1.0. The remaining genomes were assembled as follows. Single individuals were chosen per species, with preference for males when available, whose haploid status facilitates assembly. Paired-end sequencing libraries targeting 300 bp insert sizes were prepared using the Nextera protocol. Libraries were quality checked by Agilent bioanalyzer and Illumina Hi-seq sequenced to 150 base pairs (bp) by Edinburgh Genomics, United Kingdom. Sequencing for *Protobalandricus spectabilis* and additional sequencing for *Parnips nigripes* using Qiagen UltraLow Input libraries on an Illumina NextSeq mid-output 300-cycle run was performed at the ACRF Biomolecular Resource Facility, The John Curtin School of Medical Research, Australian National University. Raw reads were quality filtered and overlapping pairs merged in fastp (v0.20.1) (Chen et al. 2018) with default settings, and output fastq files were visually assessed for remaining adapters and other issues with Fastqc (v0.11.9) (Andrews 2010). Most genome assemblies were constructed using SPAdes (v3.14.0) (Bankevich et al. 2012) with most species run in isolate mode with coverage cutoff estimated automatically and default k-mers. Exceptions to this were *Cecinothofagus ibarrai, Callaspidia notata, Periclistus spJH-2016* and *Phaenoglyphis villosa*, which were assembled without a coverage cutoff and “*Aylax” hypecoi* and *Eschatocerus acacia*, which were both assembled with an additional k-mer of 99. Data for several species were first published in Hearn et al. (2019), but were re-assembled as described here for consistency (Table S1, assembly origin column). *Synergus* species genomes and the *Biorhiza pallida* and *Ganaspis species 1* transcriptomes were not re-assembled here. Quality statistics for all genomes and transcriptomes are given in Supplementary Material (Table S1).

### Gene finding

To find conserved genes suitable for phylogenetic analysis, we predicted Hymenoptera and Eukaryota BUSCOs for each genome using BUSCO v4.0.6 and OrthoDB version 10 (Simão et al. 2015) for each genome and transcriptome. The Hymenoptera dataset consisted of 5,991 BUSCO groups predicted from 40 species. Lineage-specific BUSCO datasets are composed of genes present almost universally as single copy genes, although duplications within test datasets can occur (Simão et al. 2015).

Only sequences classified as complete single copy BUSCOs were used in our analysis. A predicted BUSCO is defined as complete if its length is within two standard deviations of that BUSCO group’s mean length, that is within 95% of its expected length (Simão et al. 2015; Waterhouse et al. 2018). BUSCOs were divided into categories based on the number of species in which the gene was retrieved in a complete, single copy. In total, we found 5,890 complete single-copy BUSCOs in at least one of the 37 genomes/transcriptomes (Table 3). Our phylogenetic analyses focused on the 523 genes that were present in 34 or more of the 37 taxa (representing a total of 1.24 Mb of sequence data after alignment), and subsets of this dataset. The completeness of each genome/transcriptome assembly is given in Supplementary Material (Table S1).

**Table 3.** BUSCO gene sets and the number of species in which they were found. The total sequence length is given in nucleotide base pair equivalents; the number of amino acid sites is one third of this number.

| No. species | No. genes (cumulative) | Total length, kb (cumulative) |
| --- | --- | --- |
| 37 | 31 | 61.7 |
| 36 | 92 (123) | 224 (285) |
| 35 | 173 (296) | 350 (635) |
| 34 | 246 (542) | 602 (1,240) |
| 33 | 273 (815) | 634 (1,870) |
| 32 | 300 (1,115) | 702 (2,570) |
| 31 | 380 (1,495) | 928 (3,500) |
| 30 | 404 (1,899) | 1,020 (4,520) |
| <30 | 3,991 (5,890) | 12,200 (16,800) |

### Alignment and quality scoring

Sequences were aligned using ClustalOmega version 1.2.4 (Sievers et al. 2011), and the alignments were filtered using Gblocks version 0.91b (Talavera and Castresana 2007), with default parameters except for gap treatment, which was set to "all" to retain more phylogenetic information (Kück et al. 2010). For the purpose of phylogenetic reconstruction based on multiple genes, custom scripts were used to concatenate the desired alignments.

The putative quality of alignments was scored using the fraction of the total alignment length retained after Gblocks filtering. As alternative quality filtering and scoring options, we used HmmCleaner version 0.180750 (Di Franco et al. 2019) and OD-Seq version 1.22.0 (Jehl, Sievers & Higgins 2015), in the former case with and without previous Gblocks filtering, and in the latter after previous Gblocks filtering. HmmCleaner was used with default settings, OD-Seq with settings: distance_metric = “affine”, B = 1000, threshold = 0.025.

### Phylogenetic analysis

Phylogenetic analysis was performed using IQ-Tree version 1.6.12 (Nguyen et al. 2015) and Phylobayes version 1.8 (Lartillot & Philippe 2004) using models that accommodate site-specific amino-acid profiles. Specific settings for each program are given below.

#### IQ-Tree

We used IQ-Tree for maximum-likelihood analyses based on the C60+I+G5 model. The C60 option specifies a fast approximation of an amino-acid profile mixture model with 60 profile categories estimated from reference data (Wang et al. 2018). We modelled rate variation across sites using a mixture of invariable sites and a discrete approximation of a gamma distribution with five categories (that is, I+G5). Support values were estimated using the ultrafast bootstrap (Minh, Nguyen & von Haeseler 2013; Hoang et al. 2018) with 2,000 replicates per analysis. For each inference problem, we ran two independent analyses to confirm that phylogenetic relationships and support values were consistent. All runs used 32 CPU cores.

#### Phylobayes

In Phylobayes, we used the CAT F81 model. The CAT model infers the amino-acid profile for each site from the data assuming that the profiles come from a Dirichlet process mixture. We assumed that the exchangeability rates were the same (F81) rather than trying to estimate them from the data (the GTR option). Estimating the exchangeability rates was too computationally complex for the analyses we attempted, and it is not obvious that the results would be more accurate, as rare changes can be explained both by unusual amino-acid profiles and by low exchangeability rates under the CAT-GTR model, creating an identifiability problem that is potentially problematic. Rate variation across sites was modelled using a discrete approximation of the gamma distribution with four categories. For each inference problem, we ran two independent Phylobayes analyses for 72 hours using the MPI version on 32 CPU cores. Convergence diagnostics and consensus trees were generated for each pair of analyses using the bpcomp program in the Phylobayes package, retaining every tenth sample and using a burn-in of 25% of samples. In all cases, the mean difference in split frequencies was less than 0.005, usually much less. The maximum difference in split frequencies was 0.09 for the analysis of the problematic 36-taxon dataset (see below), but was below 0.05 for all other analyses.

#### Individual gene tree analysis

As a complement to the analyses based on concatenated gene data, we also assessed node support using metrics summarising the information for individual gene trees. We used as the species tree the one based on the best third of the alignments that include at least 34 of the 37 taxa inferred in PhyloBayes under the CAT-F81 model. Each individual gene tree was reconstructed using maximum-likelihood with the best-fit substitution model automatically selected by ModelFinder. First, using IQtree2 we calculated the gene concordance factor (gCF), which reflects the proportion of genes supporting a node considering uneven taxon sampling per gene (Minh, Hahn & Lanfear 2020). Second, using RAxML version 8.2.12, we calculated internode certainty (IC), which informs about the certainty of a bipartition by considering its occurrence in a set of gene trees relative to the occurrence of the second-best bipartition (Salichos and Rokas 2013).

### Tree figures

Illustrations of phylogenetic trees were generated using the R package ggtree version 3.2.1 (Yu et al. 2017) running under R version 4.1.1.

## Results

### Alignment quality and phylogenetic signal

We first explored the data by analyzing the dataset that contained the 31 genes that were present in all 37 taxa (Table 2). We will refer to this as the T37-G31 dataset, for 37 taxa and 31 genes. We then successively expanded the amount of data by analyzing the T36-G123 dataset (123 genes present in 36 or more taxa), T35-G296 dataset (296 genes present in 35 or more taxa) and T34-G542 datasets (542 genes present in 34 or more taxa). This series represents a trade-off between completeness in terms of taxa, and amount of genomic data included.

When these four datasets were analyzed with IQTree and a model accommodating site-specific amino-acid profiles (C60+I+G5), we discovered striking differences in topology (Fig. 1). In the smallest dataset (T37-G31; Fig. 1A), including only the complete alignments, *Eschatocerus* (Eschatocerini) diverges early in the tree. However, in the next smallest dataset (T36-G123; Fig. 1B), *Eschatocerus* is instead grouped inside the core cynipid lineages, in a clade together with *Protobalandricus* (Cynipini), *Phanacis* (Phanacidini) and *Iraella* (Aylacini s. str.). This is a somewhat surprising result, as it breaks the monophyly of the oak gall wasps (Cynipini), long presumed to be a monophyletic group. It also moves *Phanacis* (Phanacidini)—representing one major herb-galling clade—from a sister-group relationship with the other major herb-galling clade (Aulacideini) to a position within a heterogeneous collection of lineages. As more genes (and more gaps) are added, *Eschatocerus* changes again to an early-diverging position (Figs. 1C–D) but *Protobalandricus* remains outside the Cynipini, even though the support for this is quite poor in the largest dataset (Fig. 1D).

**Figure 1.**
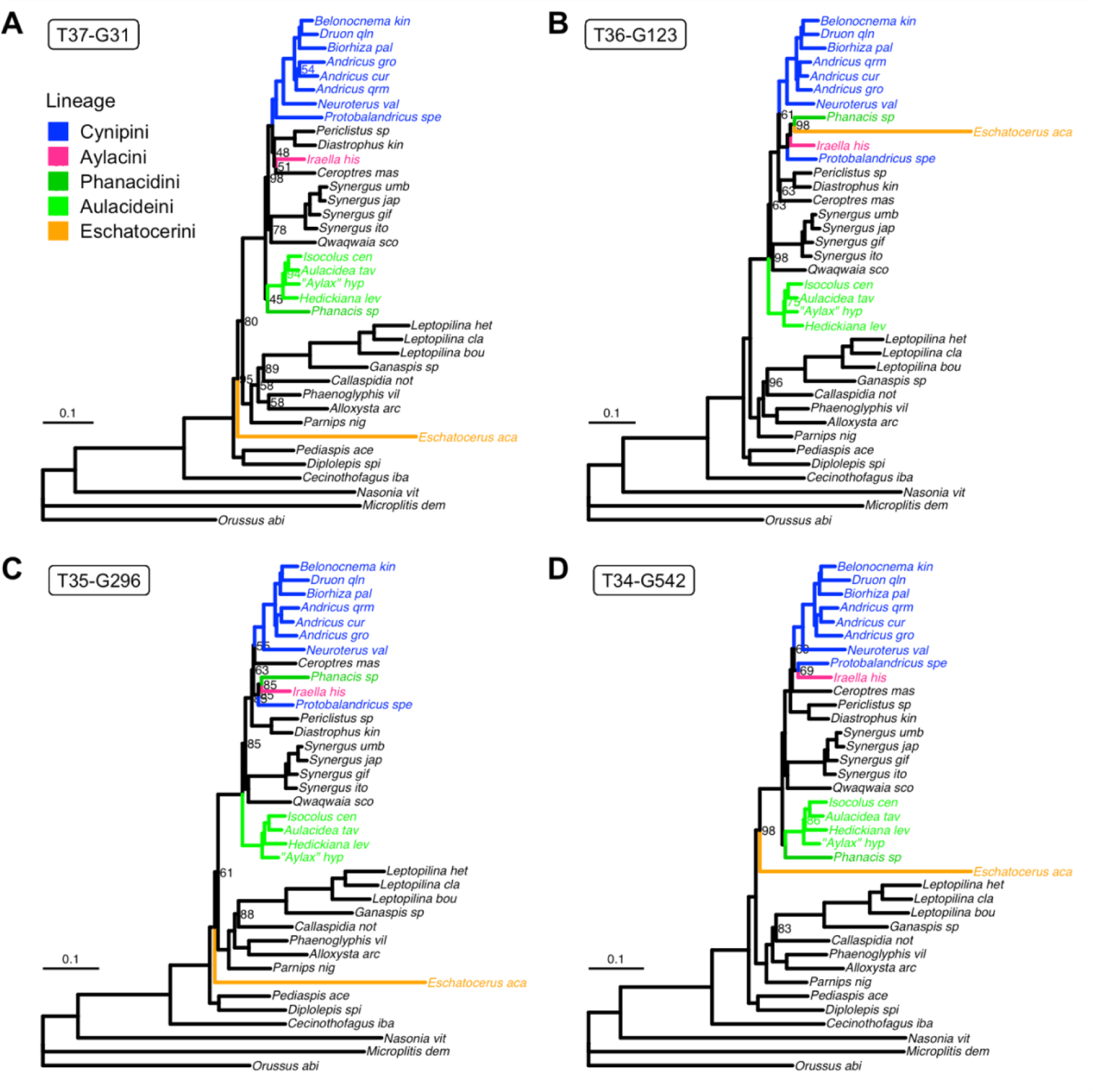
Phylogenetic relationships according to IQTree analyses of different data subsets under the C60+I+G5 substitution model. Support values (ultrafast bootstrap method) are shown on branches only if they are less than 100%. **A**. Analysis of the 31 genes present in all 37 taxa (dataset T37-G31). **B**. Analysis of the 123 genes present in 36 or more taxa (T36-G123). **C**. Analysis of the 296 genes present in 35 or more taxa (T35-G296). **D**. Analysis of the 542 genes present in 34 or more taxa (T34-G542). Note that the smallest dataset (A) results in strong support for Cynipini monophyly (blue clade). In the second smallest dataset (B), this is not the case because *Protobalandricus*, the most basal lineage in Cynipini, groups strongly with *Iraella* (Aylacini), *Phanacis* (Phanacidini) and *Eschatocerus* (Eschatocerini), the latter of which sits on a long branch. When even more data are added, the support for this assemblage successively weakens (C-D), until there is only modest evidence (69% bootstrap support) against Cynipini monophyly in the largest dataset (D).

In trying to understand these results, we noted that *Eschatocerus* is a long-branch taxon, and that the three other taxa that group with *Eschatocerus* in the next smallest dataset (Fig. 1B) have three of the five most incomplete genome assemblies in terms of the number of retrieved genes (Supplementary Material, Table S2). This suggests that the clade consisting of *Eschatocerus*, *Protobalandricus*, *Phanacis* and *Iraella* may be spurious and caused by long-branch attraction and/or poor or misleading gene alignments.

We looked at long-branch attraction first. The C60 model in IQTree is an approximation of the CAT model in PhyloBayes, and may not accurately represent site-specific amino acid profiles in cynipoids. Such deviations could potentially cause problems with long-branch attraction in our analysis. To check this possibility, we repeated the analysis of the two smallest datasets (the others were too large) in PhyloBayes using the CAT-F81 model (Fig. 2). The results were identical with those obtained with IQTree, suggesting that the topological changes are not caused by problems with the C60 approximation.

**Figure 2.**
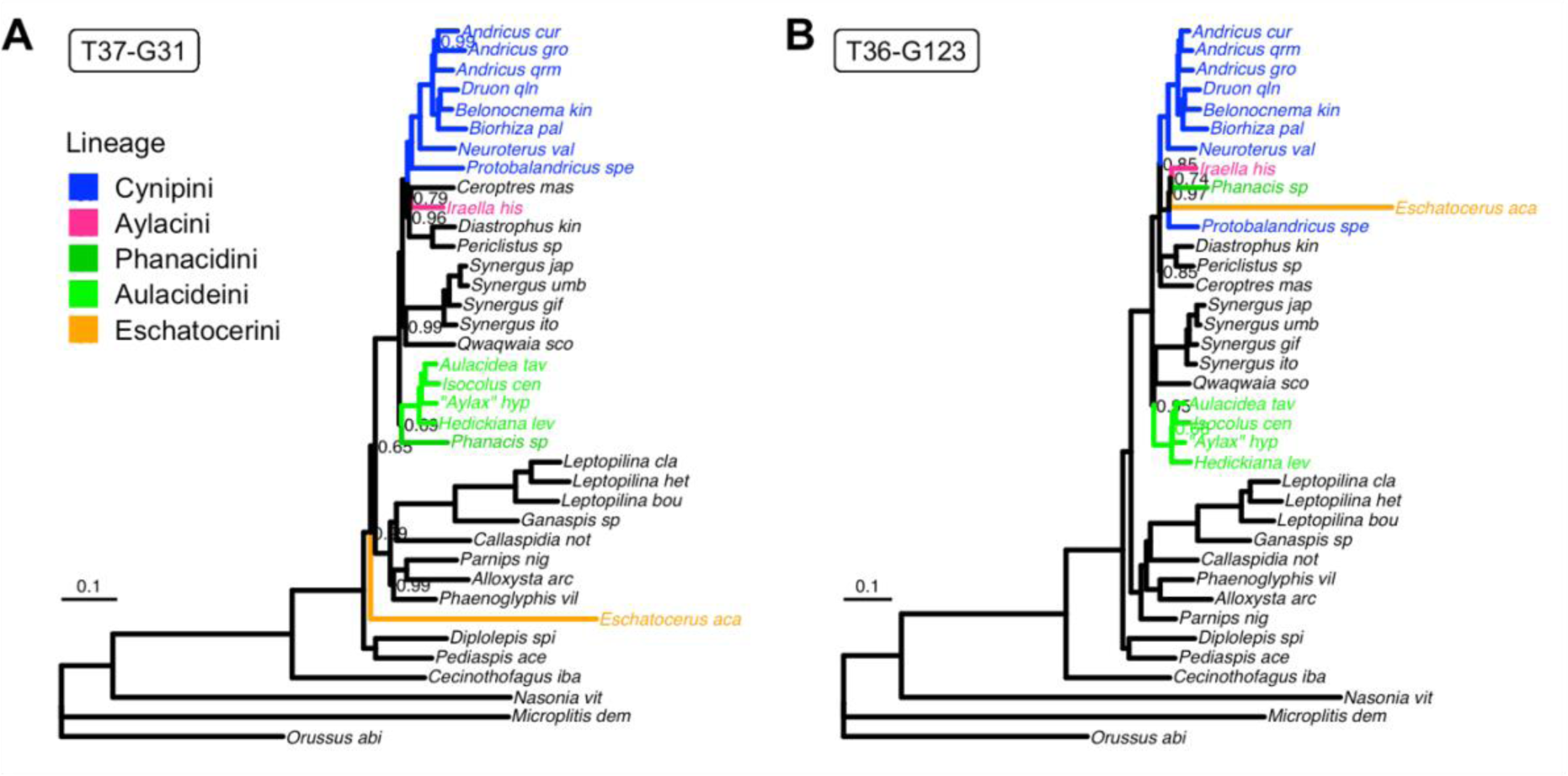
Phylogenetic relationships according to PhyloBayes analyses of the two smallest data subsets under the CAT-F81 model. Support values (posterior probability) are shown on branches only if they are less than 1.0. **A**. Analysis of the 31 genes present in all 37 taxa (dataset T37-G31). **B**. Analysis of the 123 genes present in 36 or more taxa (T36-G123). Despite the more sophisticated CAT-F81 model, which learns the amino-acid profiles from the data, the results are virtually identical to the corresponding results of IQTree (Fig. 1A, B).

Next, we turned our attention to alignment quality. We noted that even the relatively unrestrictive Gblocks filtering we used sometimes removed substantial portions of the alignments. If a substantial portion of an alignment is unreliable, then maybe also the part that remains after filtering is of doubtful quality? To examine this possibility, we divided the T34-G542 dataset into six approximately equal gene subsets based on the proportion of the alignments removed by Gblocks. When analyzed with IQTree under the C60+I+G5 model, the three best data subsets resulted in trees (Figs. 3A–C) that were identical to each other and to the tree from the no-gaps dataset T37-G31 (Fig. 1A), except for a few minor details, most of which were not well supported. Notably, *Eschatocerus* always diverged early, Cynipini was monophyletic, and *Phanacis* grouped with Aulacideini in all these trees.

**Figure 3.**
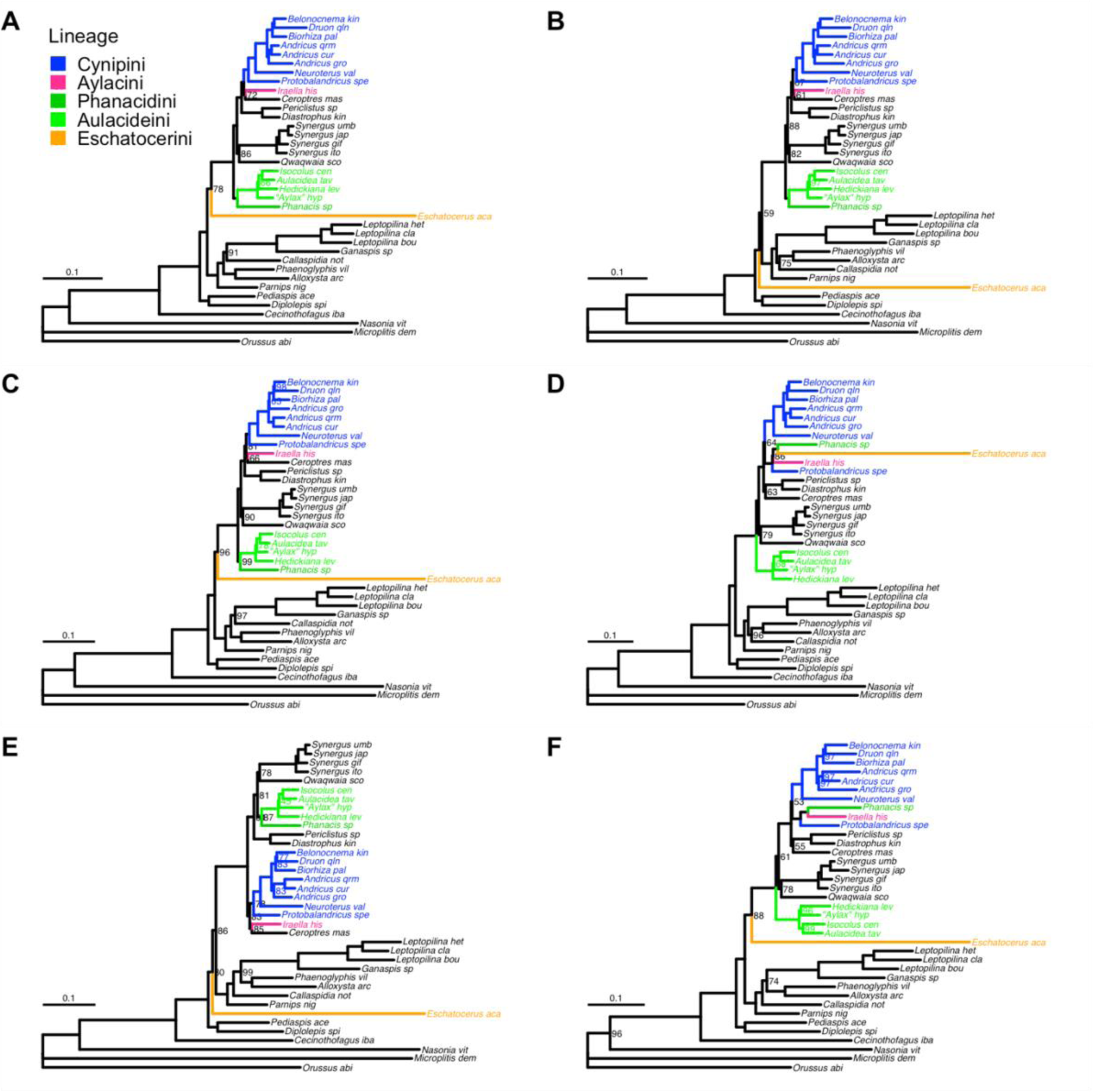
Phylogenetic results for six equally-sized, quality-ranked subsets of the T34-G542 dataset, analyzed using IQTree under the C60+I+G5 model. The raw alignments were subjected to filtering and quality ranking by Gblocks. Support values (ultrafast bootstrap) are shown on branches only if they are less than 100%. **A**. Less than 13% of sites filtered out (best quality). **B**. From 13% to 26% filtered out. **C**. From 26% to 37% filtered out. **D**. From 37% to 47% filtered out. **E**. From 47% to 59% filtered out. **F**. More than 59% filtered out (worst quality). The three best subsets (A-C) yield congruent results except for the position of *Eschatocerus*, which varies slightly but without strong conflict in support values. All have monophyletic Cynipini (blue lineages), and none of them group Phanacidini, Aylacini and Eschatocerini with each other or with *Protobalandricus*, as seen in some of the poor-quality data subsets (D, F).

The results for the three worst data subsets (Figs. 3D–F) differed among themselves and from the results of the no-gaps dataset (T37-G31) in several respects, often involving aberrant placements of the four problematic taxa mentioned previously—*Eschatocerus*, *Protobalandricus*, *Phanacis* and *Iraella*—or unusual arrangements of more basal branching events, but with low support. Thus, it appears that the phylogenetic signal is consistent in the best gene alignments, that is, those that contain only small portions that are detected as problematic by the Gblocks filter.

To further test the effect of alignment quality, we also explored partitions of the T34-G542 dataset generated using other filtering and scoring methods. Specifically, we tried HmmCleaner, Gblocks + HmmCleaner and Gblocks + OD-Seq, and then divided the gene alignments into subsets based on how many sites were removed (HmmCleaner and Gblocks + HmmCleaner) or how many sequences (Gblock + OD-Seq) were removed by each of these pipelines. In all cases, the IQTree analyses of the highest-quality data subset or subsets resulted in trees that were identical or almost identical to the tree from the no-gaps analysis (Supplementary Material, Figs. S1–S3). OD-Seq improves the quality of alignments by removing sequences that appear to represent outliers. Thus, the quality scores are few, and it was difficult to devise criteria that generated partitions of equal size. We therefore ended up with the best partition (no sequences removed) being much smaller than the other ones (approximately 2,900 sites versus 25,600–56,800 sites), and resulting in a poorly resolved tree with some unusual features (Supplementary Material, Fig. S3A). Analysis of the next best OD-Seq partition, however, retrieved a tree that was highly similar to the no-gaps tree (compare Fig. 1A to Supplementary Material, Fig. S3B).

Based on these results, we conclude that it is mainly poor-quality alignments that generate the somewhat unexpected placements of *Eschatocerus*, *Protobalandricus*, *Phanacis* and *Iraella* in analyses of the T36-G123, T35-G296 and T34-G542 datasets.

### Long-branch attraction and gene tree discordance

The tree on which all analyses of high-quality alignments converge (e.g., Figs. 3A–C) supports many previous notions of cynipid relationships. For instance, the cynipid tribes Cynipini, Diastrophini, Synergini (s. str.) and Aulacideini are all monophyletic, as is the family Figitidae, the figitid subfamilies Eucoilinae and Charipinae, and the Cynipoidea as a whole. However, somewhat surprisingly, the gall wasps themselves (Cynipidae) are not monophyletic. The Diplolepidini + Pediaspidini (represented by *Diplolepis* and *Pediaspis*) and Paraulacini (represented by *Cecinothofagus*) lineages both fall outside a core cynipid clade that apparently constitutes the sister group of the Figitidae. In some analyses, the Eschatocerini (represented by *Eschatocerus*) also fall outside this clade.

The putative cynipid lineages that place outside the core cynipid clade, however, all represent long branches in the tree, as do the outgroup taxa. Could the non-monophyly of Cynipidae be the result of long-branch attraction, pulling isolated cynipid lineages towards the outgroups? To examine this question, we focused on a dataset consisting of the two best subsets of the T34-G542 dataset according to the Gblocks criterion, and we used PhyloBayes for the best chances of detecting long-branch attraction. The analysis of the complete taxon set resulted in the tree with non-monophyletic Cynipidae (Fig. 4A). From this dataset, we then removed in turn *Cecinothofagus*, *Eschatocerus*, outgroups, *Cecinothofagus* + *Eschatocerus*, and *Eschatocerus* + outgroups. These were the only removals of long-branch taxa that left a sufficient number of remaining lineages to test non-monophyly of Cynipidae. In all cases, the support for non-monophyletic Cynipidae remained at 100% (Figs. 4B–F). The results were almost identical when the same datasets were analyzed with IQTree (Supplementary Material, Fig. S4).

**Figure 4.**
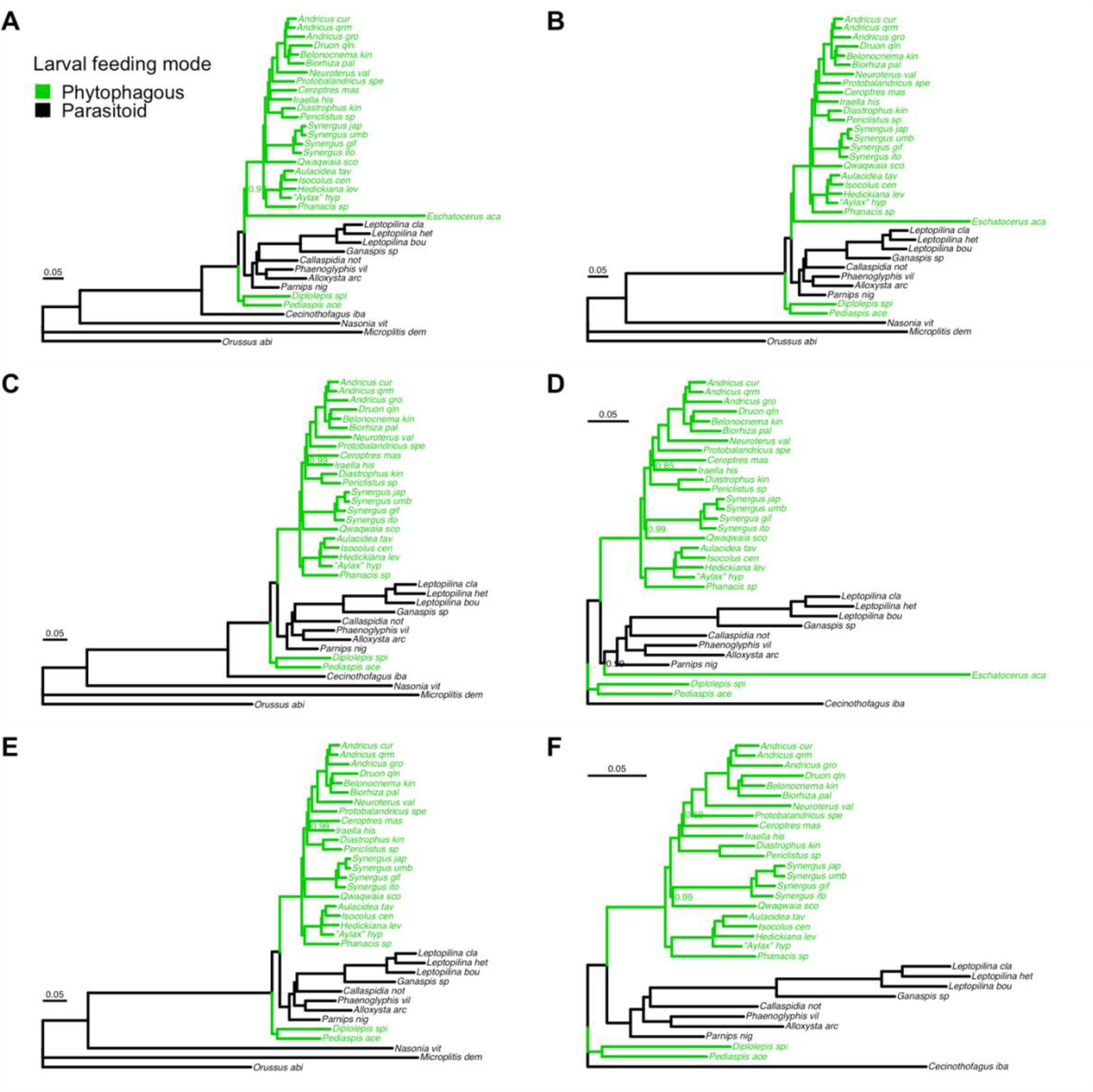
Testing the potential effect of long-branch taxa on phylogenetic results. For these analyses, we used the best third of the T34-G542 alignments, that is, the alignments where Gblocks filtered out 26% or less of the sites (see Fig. 3). For the best possibility of detecting model-related long-branch attraction effects, we used PhyloBayes and the CAT-F81 model. Branch support values (posterior probability) are only shown if they are less than 1.0. **A**. Analysis of the full taxon set. **B**. *Cecinothofagus* excluded. **C**. *Eschatocerus* excluded. **D**. Outgroups excluded. **E**. *Cecinothofagus* and *Eschatocerus* excluded. **F**. *Eschatocerus* and outgroups excluded. Regardless of taxon exclusion, the relationships among the included lineages remain identical to those in the full analyses, except for a slight variation in the position of *Eschatocerus* when outgroups are excluded (D). Notably, the phytophagous groups (gall inducers and inquilines, green) remain diphyletic in all analyses with respect to the parasitoid lineages (black).

The gene tree concordance analysis shows that there is consistent signal across gene trees for the deep splits in the superfamily, that is, between *Cecinothofagus* and the remaining taxa, and between *Diplolepis* + *Pediaspis* on one hand and the remaining Cynipidae and Figitidae on the other (Supplementary Material, Fig. S5). This is reflected both by a positive internode certainty and a gene concordance factor > 40%. The relationships among *Eschatocerus*, remaining Cynipidae (s. str.) and Figitidae are not consistently resolved across gene trees. Similarly, this analysis indicates a fair amount of inconsistency across gene trees concerning tribal relationships within the Cynipidae (s. str.) excluding *Eschatocerus*. This could be because errors in the assemblies, errors in gene tree inference due to lack of data or biases in the simplified model used, or true inconsistencies among the gene trees. However, we did not pursue this further.

### Phylogenetic relationships

As our best phylogenomic estimate of relationships, we present the PhyloBayes (CAT-F81) analysis of the two best subsets of the T34-G542 dataset according to the Gblocks criterion (Fig. 5; see also Fig. 4A and Supplementary Material, Fig. S4A). On the tree, we have indicated the currently recognized cynipid tribes, and a proposed reclassification of the family Cynipidae into three family-level taxa: the Cynipidae (s. str.) for the core cynipid clade, including Eschatocerini; the Diplolepididae for Diplolepidini + Pediaspidini; and the Paraulacidae for the Paraulacini.

**Figure 5.**
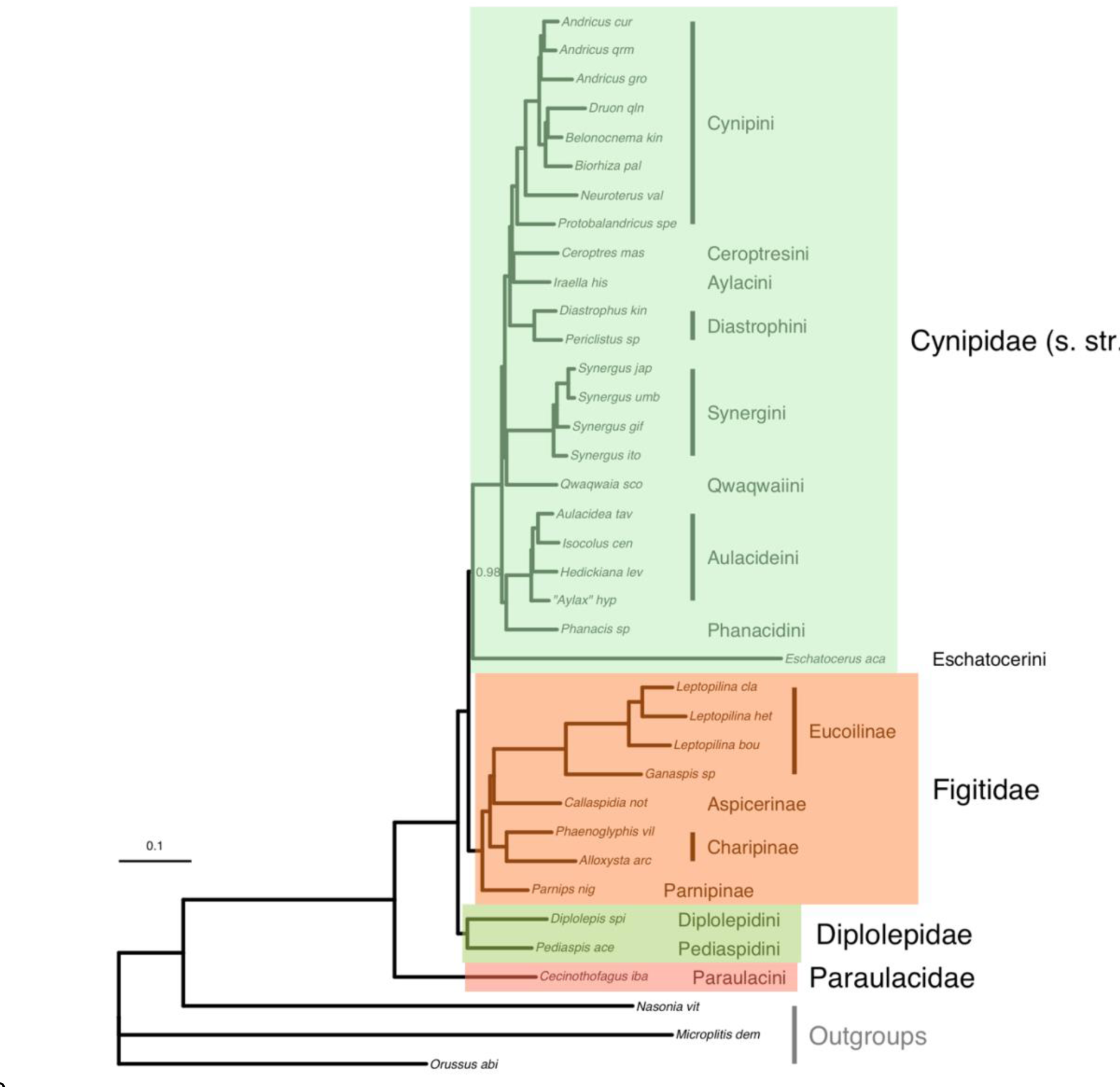
Preferred hypothesis of phylogenetic relationships. The tree is based on the best third of the alignments that include at least 34 of the 37 taxa (T34-G542 dataset, Gblocks filtering removed less than 26% of sites), analyzed using PhyloBayes under the CAT-F81 model (the same analysis shown in Fig. 4A). Current cynipid tribes and figitid subfamilies are indicated, together with the proposed new classification of cynipid lineages into three distinct families.

Our results suggest that the two major tribes of herb gallers, Phanacidini and Aulacideini, form a natural group at the base of the Cynipidae (s. str.). The third tribe of herb gallers (Aylacini (s. str.)), represented in our analysis by *Iraella*, is apparently more closely related to the oak gallers (Cynipini) and the oak inquilines in the tribe Ceroptresini (represented by *Ceroptres*) than to the other herb gallers. The Aylacini (s. str.) are all associated with plants in the family Papaveracae. The Phanacidini and Aulacideini are most commonly associated with Asteraceae and Lamiaceae but there is one species in the Aulacideini associated with Papaveraceae, “*Aylax*” *hypecoi*. This species was included in our analysis, and our results confirm that this species is not a member of Aylacini (s. str.), consistent with previous analyses (Ronquist et al. 2015).

The Diastrophini, represented in our analysis by *Periclistus* and *Diastrophus*, form the sister group of the clade including Cynipini + Ceroptresini + Aylacini (s. str.). It is a tribe that includes both inquilines and gall inducers associated with host plants in the family Rosaceae, mostly bushes of the genera *Rubus* and *Rosa* but also herbs in the genus *Potentilla*.

The tribe Qwaqwaiini, represented in our analysis by the only described species, *Qwaqwaia scolopiae*, appears to be the sister group of the clade formed by Synergini (s. str.) + Rhoophilini, which mostly includes inquilines in Cynipini galls and a few other insect galls. However, one of the species we analyzed, *Synergus itoensis*, represents a small subgroup within the Synergini (s. str.) of true gall inducers on oaks. This subgroup of gall inducers appears to be the sister group of the rest of the Synergini in our analysis only because several early-diverging representatives are missing (Ronquist et al. 2015; Ide et al. 2018; Lobato-Vila et al. 2022). The Qwaqwaiini + (Synergini (s. str.) + Rhoophilini) apparently represent the sister group of the remaining Cynipidae (s. str.), except for the Eschatocerini. The latter tribe, represented in our analysis by the single genus *Eschatocerus*, appears to be the sister group of all other Cynipidae (s. str.). Occasionally, we retrieved *Eschatocerus* as the sister group of remaining Cynipidae (s. str.) + Figitidae (Fig. 3B, Supplementary Material, Figs. S2A–B, S3B) or only Figitidae (Supplementary Material, Fig. S1B), although with unconvincing support values. Thus, we conclude that the tribe Eschatocerini likely belongs to the Cynipidae (s. str.).

The Figitidae in our analyses form a strongly supported monophyletic group. The subfamily Parnipinae, represented in our analysis by the single genus *Parnips*, appears as the sister group of the remaining lineages. It is a parasitoid of cynipid gall inducers in the tribe Aylacini (s. str.). The Charipinae, represented by *Phaenoglyphis* and *Alloxysta* in our analysis, form a monophyletic group. They are hyperparasitoids of other parasitic wasps attacking aphids. The remaining Figitidae apparently form a monophyletic group, falling into two subgroups: the Aspicerinae (Callaspidia) and the Eucoilinae (the remaining species). Both subfamilies are parasitoids of Diptera larvae.

Among the more early-diverging cynipoid lineages, the Diplolepidini, represented by *Diplolepis*, and the Pediaspidini, represented by *Pediaspis*, form a strongly supported clade, which appears to be the sister group of Figitidae + Cynipidae (s. str.). We propose here that this clade be recognized as a separate family, the Diplolepididae (Fig. 5).

Finally, our results support the conclusion that the Paraulacini (represented by *Cecinothofagus*) form the sister group of the remaining cynipoid lineages. We propose here that also this clade be recognized as a separate family, the Paraulacidae (Fig. 5). The new classification is discussed in more detail in the Taxonomy section below.

### Evolutionary implications

Our analysis includes too few exemplars to allow a rigorous statistical analysis of different scenarios for the origin of cynipoid gall inducers and inquilines, but we illustrate some of the possibilities (Figs. 6, 7). When it comes to the origin of the phytophagous lineages (inquilines and gall inducers), two main alternative scenarios seem plausible.

**Figure 6.**
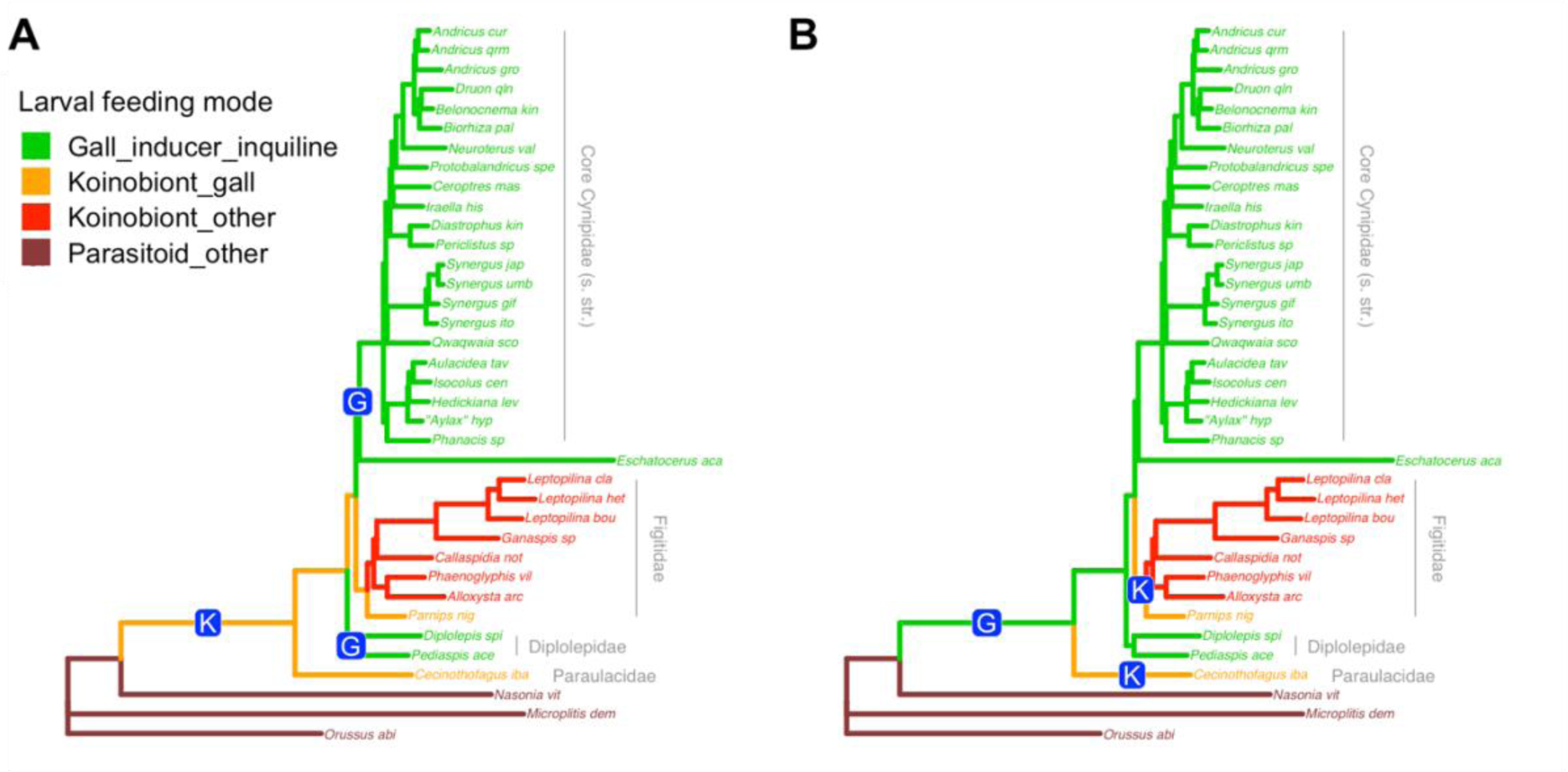
Two possible scenarios for the origin of major life-history types in the Cynipoidea. **A**. *Independent phytophagy scenario*. The ancestor of cynipoids was a koinobiont endoparasitoid (at least in early instars) of gall inducing insects (orange lineages, origin of koinobionts marked with “K”). The ancestral life history persists today in the Paraulacidae and basal lineages of Figitidae, like the Parnipinae. Gall inducers and inquilines originated twice from these koinobionts of gall insects (“G”). **B**. *Parasitoid reversal scenario*. The koinobiont endoparasitoids of gall insects (“K”) evolved independently in the Paraulacidae and Figitidae, possibly in both cases from phytophagous gall inducers and inquilines (“G”). In both scenarios, advanced figitid lineages (in red) remained koinobiont parasitoids of insects but colonised hosts in other environments.

**Figure 7.**
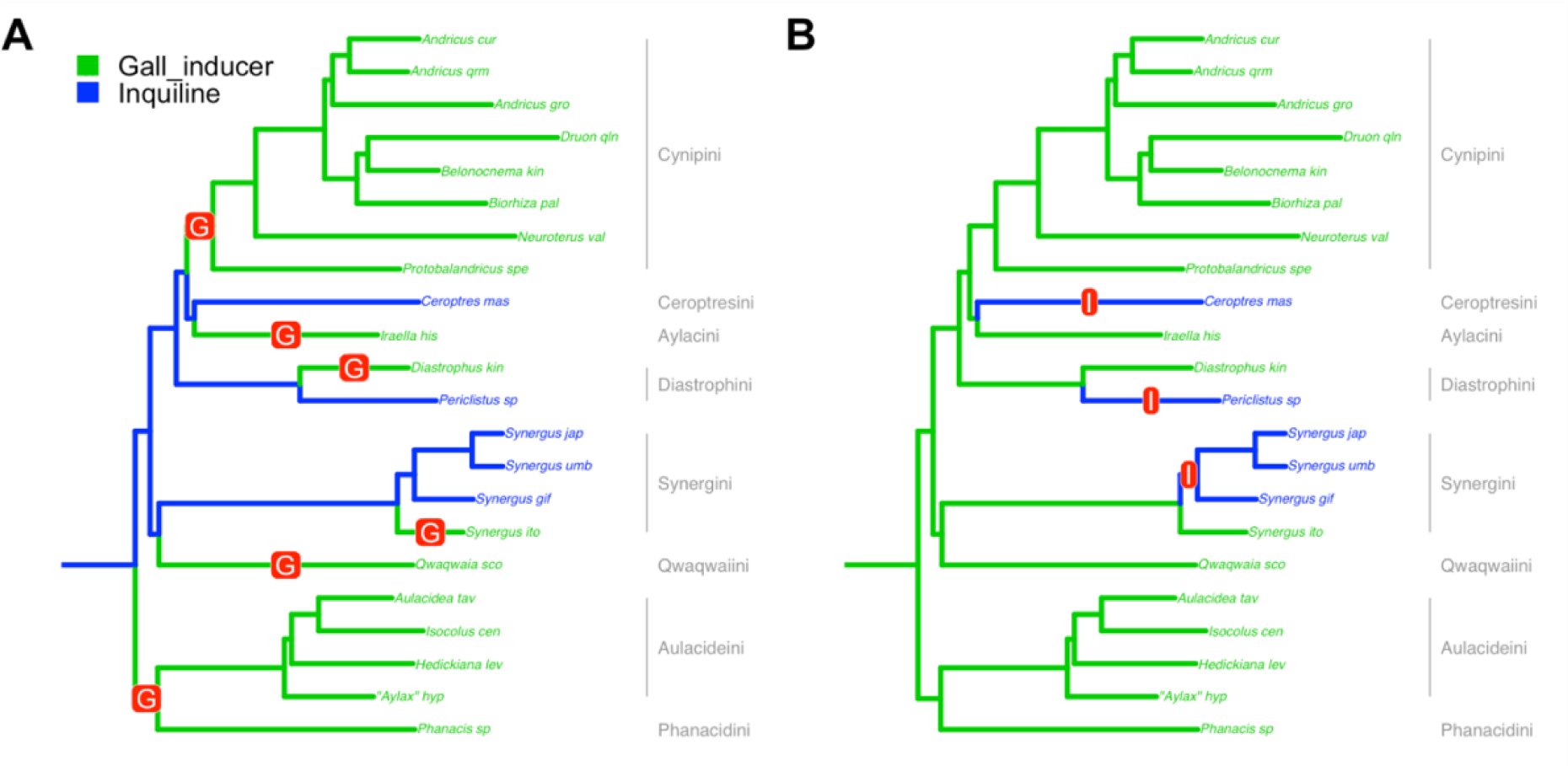
Two possible extreme scenarios for the origin of gall inducers (green lineages) and inquilines (blue lineages) in the Cynipidae (s. str.) (*Eschatocerus* excluded because of uncertainty concerning its life history). **A**. *Inquilines-first scenario*. Gall inducers evolved repeatedly from inquilines, which represent an intermediate stage in the origin of true gall inducers. At least six independent origins of gall inducers will have to be assumed for the included lineages, seven if additional evidence is considered (see text). **B**. *Gallers-first scenario*. In this scenario, inquilines represent gall inducers that have lost the ability to initiate galls. At least three independent origins of inquilines will have to be assumed for the included lineages, ten if additional evidence is considered (see text). In reality, evolution may have taken a path that involved transitions in both directions.

One scenario (independent phytophagy) assumes that the Paraulacidae and Parnipinae life histories trace back to the ancestral cynipoid (Fig. 6A). If so, then phytophagous inquilines or gall inducers must have originated twice from such ancestors. The other scenario (parasitoid reversal) assumes instead that it is the phytophagous habit that traces back to the cynipoid ancestor (Fig. 6B). Then koinobiont endoparasitoids must have evolved twice independently from phytophagous ancestors. Intermediates between these extremes are also possible; for instance, gall inducers and koinobiont endoparasitoids might both have evolved twice independently from ancestors that were ectoparasitoids of gall-inducing insects.

Inferring the evolutionary transitions between inquilines and gall inducers is even more complicated. We present two extreme scenarios. If all gall inducers evolved from inquilines (inquilines-first scenario), then our results show that gall inducers must have evolved at least six times independently (Fig. 7A). If additional evidence on the phylogeny of Diastrophini (Ronquist et al. 2015) is taken into account, this increases to seven. At the other extreme, if all inquilines evolved from gall inducers (gallers-first scenario) our results taken at face value suggest three transitions (Fig. 7B). However, the gall-inducing *Synergus* lineage is deeply nested within inquiline lineages (Ide et al. 2018; Lobato-Vila et al. 2022), and several additional, independent origins of inquilines would be required if the gall-inducing habit of this lineage is ancestral. If we also consider inquilines in the Diastrophini (Ronquist et al. 2015), then there would have been at least ten independent origins of inquiline cynipids from gall inducers. Of course, intermediate scenarios involving transitions in both directions are possible, even though there is no simple solution with only few switches between the two life histories.

### Taxonomy

#### Taxonomy

Given that our results provide solid and independent confirmation of the results from the UCE analysis (Blaimer et al. 2020) regarding the non-monophyly of Cynipidae, we find it appropriate to revise the family-level classification to reflect these findings here. As the circumscription of the 13 cynipid tribes and potential apomorphies characterizing each of them have been discussed at length previously (Ronquist et al. 2015; Lobato-Vila et al. 2022), we just give a brief formal synopsis of the proposed family classification here. The synopsis does not include the fossil family-level taxa, the phylogenetic position of which must be carefully re-evaluated in light of the phylogenomic findings. We refrain from revising the classification of the non-cynipid family-level taxa in the Cynipoidea, as the results of the UCE analysis on Liopteridae, Ibaliidae and some Figitidae lineages still await independent confirmation.

#### Paraulacidae Nieves-Aldrey and Liljeblad, stat. prom

[ZooBank identifier to be inserted upon acceptance] Type genus *Paraulax* Kieffer, 1904

Paraulacini Nieves-Aldrey and Liljeblad, 2009.

#### Circumscription

The family includes the genera *Paraulax* and *Cecinothofagus,* each with three species. Southern South America, found only in the temperate *Nothofagus* forests of Chile and Argentina.

#### Comments

A set of unique morphological features allow easy differentiation of Paraulacidae from Cynipidae and other families in Cynipoidea (Ronquist et al. 2015). Two unique autapomorphies can be emphasized: the presence of 5-9 vertical carinae in the ventral region of the gena; and the profemur with the basal third swollen and carrying a structure of 4-5 rows of sharp, closely spaced and deep costulae. The Paraulacidae appear to be parasitoids of gall inducing chalcidoids of the genus *Aditrochus* on species of *Nothofagus* (Nothofagaceae) (Rasplus, Nieves-Aldrey & Cruaud 2022).

#### Diplolepididae Latreille, 1802, stat. prom

[ZooBank identifier to be inserted upon acceptance]. Type genus *Diplolepis* Geoffroy, 1762. Conserved (see Kerzhner 1991).

Diplolepariae Latreille, 1802. Corrected to Diplolepididae. Rhoditini Hartig, 1840.

Diplolepidini Latreille (Ronquist 1999)

#### Circumscription

Includes two tribes, Diplolepidini and Pediaspidini.

Diplolepidini Latreille, 1802. Two genera, *Diplolepis* Geoffroy with 51 species and *Liebelia* Kieffer with 10 species. Holarctic.

Pediaspidini Ashmead, 1903. Two genera, *Pediaspis* and *Himalocynips*, with one species each. Palaearctic.

### Comments

*Himalocynips*, a genus with a single species that was described within its own subfamily (Himalocynipinae Yoshimoto, 1970) was included within the Pediaspidini by Ronquist (1999). Its phylogenetic proximity to *Pediaspis* was later supported by a morphological phylogenetic analysis (Liljeblad et al. 2008). The biology and host plant of this species are however unknown, although it may (as for *Pediaspis*) be a galler on *Acer* (Sapindaceae). We were unable to include this rare and poorly studied species in our analysis, and a molecular confirmation of its placement within the Pediaspidini and the Diplolepididae is an obvious priority for future studies.

Putative morphological apomorphies for the Diplolepidini include the ploughshare-shaped hypopygium, the broad and crenulate mesopleural impression, and the lack of lateral propodeal carinae (Ronquist et al. 2015). However, quantitative analyses have struggled to identify unique or distinct apomorphies for the tribe, partly because of variation among the constituent taxa, and partly because of the previous difficulties in resolving relationships among cynipid tribes (Ronquist et al. 2015). The Pediaspidini are characterized by several unique or distinct apomorphies, among them the posteromedian scutellar impression (Ronquist et al. 2015). Potential apomorphies of the Diplolepididae include the faint or absent scutellar foveae and the female antenna having 12 or more flagellomeres (Ronquist et al. 2015; couplet 3 in the key to cynipid tribes). A quantitative analysis of the available morphological and biological evidence for Diplolepididae monophyly is still missing. Before such an analysis is attempted, however, it would be valuable to reassess the morphological evidence in the light of the phylogenomic results. It is clear that such an analysis would have to span all major cynipoid lineages, and not be restricted to the former cynipid groups.

We refrain from elevating the Diplolepidini and Pediaspidini to subfamily status, as we think it is likely that further study of the Eastern Palaearctic fauna will reveal additional divergent lineages within the family.

### Cynipidae (s. str.)

Cynipsera Latreille, 1802. Corrected to Cynipidae. Type genus: *Cynips*.

### Circumscription

As here proposed, the family includes the tribes Eschatocerini, Phanacidini, Aulacideini, Qwaqwaiini, Synergini, Rhoophilini, Diastrophini, Aylacini, Ceroptresini and Cynipini.

### Comments

The position of the Eschatocerini is still highly uncertain, and its life history is also poorly studied. Future studies will have to show whether it truly belongs to the Cynipidae (s. str.), or elsewhere in the Cynipoidea, probably then as a separate family. The potential apomorphies of each of the remaining tribes have been analyzed previously (Ronquist et al. 2015), although it would be valuable to reassess the morphological and biological evidence and reanalyze it in the light of the new phylogenomic results. The same applies to potential apomorphies for the Cynipidae (s. str.). In the latter case, there are no known apomorphies at present.

Although there is growing evidence that the Phanacidini and Aulacideini are sister groups, we prefer to keep them as separate tribes (in contrast to Blaimer et al. 2020), as there are distinct biological and morphological differences between the groups. The Phanacidini tend to be small and elongate species, and most of them are stem gallers. The Aulacideini tend to be larger and their body form is more rounded. They usually induce galls in other plant parts.

Although one could argue for the grouping of tribes within the Cynipidae (s. str.) into subfamilies, we consider it premature to do so at the current time. In particular, it would be advantageous if the position of the Eschatocerini could be determined unambiguously before further refinement of the classification is considered.

## Discussion

### Alignment quality and phylogenetic signal

Assembling genomes or transcriptomes from short sequence reads and finding single-copy orthologs in those assemblies are challenging tasks. Thus, one might expect some variation in the quality of the resulting gene datasets. There is a plethora of tools for aligning the gene sequences in those datasets, and for filtering out alignment sites or sequences that may provide noisy or misleading phylogenetic signal. Nevertheless, it may be difficult to eliminate such data issues. Our phylogenetic results varied depending on which gene alignments were included but were consistent for the high-quality alignments, regardless of method used to identify the latter (alignment completeness, Gblocks results, HmmCleaner results, Gblocks + HmmCleaner results, or Gblocks + Od-Seq results).

This suggests that we had problems with misleading phylogenetic signal in poor alignments, rather than true conflict between different gene trees. This is also supported by the fact that the four taxa that were apparently incorrectly grouped together in analyses including poor alignments (*Eschatocerus, Iraella, Phanacis* and *Protobalandricus*) also were represented by some of the most incomplete genome assemblies. It is interesting to note that the taxa represented by transcriptomes (*Biorhiza* and *Ganaspis*) were not affected by similar problems with unstable phylogenetic placements, despite the rather incomplete representation of the genome in these transcriptomes (Supplementary Material, Table S2). This, too, supports the conclusion that some alignments included misleading phylogenetic signal from poor-quality genome assemblies, and gives some confidence in the tree resulting from analysis of the high-quality alignments. Interestingly, our results also suggest that quality filtering tools, such as the ones we tested (Gblocks, HmmCleaner and OD-seq), are better at identifying problem alignments than they are at filtering out erroneous or misleading sites and sequences. None of these tools were able to remove the misleading phylogenetic signal from the poor alignments, although they might have had some positive effect.

The ultimate cause of the discordant phylogenetic signal remains unclear. The four problematic taxa may group together in some analyses simply because they share divergent or incorrect sequences for some genes. The signal could be entirely erroneous - for example through sharing of specific gene pairs that can easily be merged into chimeric sequences in challenging genome assemblies, resulting in positively misleading phylogenetic signal that groups them together.

As several alternative approaches to filtering out poor gene alignments gave consistent end results, we are fairly confident that our phylogenetic analysis is not misled by erroneous genome assemblies. It is more difficult to exclude the possibility of shortcomings in the substitution model used for probabilistic inference resulting in artificial long branch attraction. Resolving cynipoid relationships involves determining the branching order of several long branch taxa (i.e., groups linked to other members of the taxon set by a long, non-dividing branch inserting deep in the phylogeny), including the Eschatocerini, Paraulacidae and Diplolepididae. The problem is accentuated by the long evolutionary distance between known cynipoid and outgroup genomes. By using models that accommodate among-site variation in amino-acid profiles, we applied some of the best available tools for resolving long-branch attraction due to model shortcomings (Kapli and Telford 2020). We also note that removal of long-branch taxa in various combinations revealed no sign of an alternative phylogenetic signal obscured by long-branch attraction effects (Fig. 4).

### Phylogenetic relationships

The phylogenetic results from our analysis are largely congruent with and complement those of the earlier UCE analysis (Blaimer et al. 2020). Here we highlight the major agreements and disagreements between these two phylogenomic studies.

i. *Division of Cynipidae into 3 families and placement of Eschatocerini*. Both studies support division of the Cynipidae into three separate lineages—recognized here as the families Paraulacidae, Diplolepididae and Cynipidae (s. str.). However, the evidence on the placement of Eschatocerini is slightly different. Our analysis suggests that the Eschatocerini belong to the Cynipidae (s. str), forming the sister-group of the remaining lineages in that clade, while the UCE analysis favored a sister-group relationship between the Eschatocerini and Figitidae. As the *Eschatocerus* genome assembly is one of the least complete in our study, further genomic sequencing of this taxon would be highly desirable.
ii. *Rejection of monophyly of cynipid herb gallers*. Our study provides even stronger support for the conclusion of the UCE analysis (Blaimer et al. 2020) that the herb-galling clade of Aulacideini + Phanacidini is monophyletic. Both analyses are consistent with previous studies suggesting that these two tribes are reciprocally monophyletic (Liljeblad & Ronquist 1998; Ronquist et al. 2015). Unlike Blaimer et al. (2020), we prefer to keep the tribes Aulacideini and Phanacidini separate until there is evidence that this would conflict with phylogenetic relationships. Blaimer et al. (2020) also concluded that the cynipid herb gallers (apart from a few species in the tribe Diastrophini) form a monophyletic clade, Aylacini (s. lat.), which is the sister group of all remaining Cynipidae (s. str.). As mentioned above, this interpretation is based on the incorrect assumption that *Neaylax salviae* (which they name *Aylax salviae*) belongs to the Aylacini (s. str.). In fact, this species belongs to a clade of Lamiaceae gallers in the Aulacideini (Ronquist et al. 2015), and is unrelated to the true Aylacini (s. str.), all known species of which are associated with poppies (Papaveraceae). Our study is the first phylogenomic analysis to include a true representative of the Aylacini (s. str.), *Iraella hispanica*, and our analysis clearly shows that herb gallers in Aylacini (s. str.) and in Aulacideini + Phanacidini are phylogenetically divergent. Instead, Aylacini (s. str.) is deeply nested within the sister-group of Aulacideini + Phanacidini, a clade that is dominated by inquilines and gall inducers associated with woody rosids (the only exception being a few species of Diastrophini that are gallers of herbs in the genus *Potentilla*). Thus, galling of herbs in the family Papaveraceae by the Aylacini (s. str.) appears to be secondary. Our results are consistent with several earlier analyses suggesting that the Aylacini (s. str.) form a lineage that is distinct from that of the Aulacideini and Phanacidini (Liljeblad & Ronquist 1998; Nylander et al. 2004; Ronquist et al. 2015), and they agree with preliminary analyses of a recent genome assembly of *Aylax minor*, another member of the Aylacini (s. str.) (AB, unpublished data).
iii. *Phylogenetic placement of the Qwaqwaiini*. Ours is the first phylogenomic analysis to include the Qwaqwaiini. Our analysis places *Qwaqwaiia scolopiae*, the only known species in the Qwaqwaiini, as the sister group of the inquiline clade consisting of Synergini (s. str.) + Rhoophilini. This is intriguing because, like the Qwaqwaiini, the Rhoophilini are from South Africa. To date these are the only two indigenous species of Cynipidae (s. str.) known from the afrotropical zone. This could potentially suggest an afrotropical origin for this clade.
iv. *Relationships in Figitidae*. Our sampling of Figitidae is not as extensive as Blaimer et al.’s UCE analysis, but our results are entirely consistent for all taxa that overlap. In both analyses, the Parnipinae is the sister-group to all other Figitidae, the Charipiniae (*Phaenoglyphis* and *Alloxysta* in our analysis) are monophyletic, the Diptera-associated lineages (Aspicerinae (*Callaspidia*) and Eucoilinae (*Ganaspis* and *Leptopilina*) in our analysis) form a monophyletic group, and the Eucoilinae are monophyletic. As our analysis did not include any representatives of Liopteridae and Ibaliidae, we cannot comment on their placement. Neither our analysis nor any other molecular phylogenetic analysis has yet included representatives of the rare Australian Austrocynipidae, which is assumed to be the sister group of all other cynipoids (Ronquist 1995, 1999).

### Evolutionary implications

#### Transitions between phytophagous and parasitoid lifecycles

The phylogenomic results suggest two alternative scenarios for the origin of gall inducers and inquilines from insect-parasitic ancestors (Fig. 6). Superficially, it may appear more likely that the phytophagous forms evolved once, and that figitids secondarily reverted to an insect-parasitic life history (Fig. 6B). If so, and given that Paraulacidae are koinobiont endoparasitoids of gall-inducing insects, then adaptations to this specialized mode of parasitism must have evolved separately in the Paraulacidae and Figitidae. The alternative hypothesis requires independent origins of herbivory (inquilinism/gall-induction) in the Diplolepididae and Cynipidae (s. str.) from parasitoids of gall insects (Fig. 6A). Assessing which of these patterns of transition is most probable requires that we weight the relative probabilities of the alternative state changes. Such weighting requires more information than currently available.

Interestingly, both Diplolepididae and Cynipidae include species whose genomes encode plant cell wall degrading enzyme genes (Hearn et al. 2019). These may have been acquired from an herbivorous shared common ancestor, or alternatively they may be essential components of cynipid herbivory that have been acquired convergently during independent evolution of galling lifestyles. Analyses of whether such complex genomic features associated with the two different life histories are likely to have a shared history or separate origins provides one of the most promising ways of distinguishing between the two possible scenarios.

Discrimination between the alternative hypotheses shown in Figs. 6A and B is made more difficult by uncertainty regarding the biologies of some of the taxa involved. It would be good to have additional data confirming that the Paraulacidae are indeed koinobiont endoparasitoids, and identifying the Eschatocerini as true gall inducers, inquilines or parasitoids. Demonstration of herbivory for *Eschatocerus* (as we have assumed in Fig. 6) would strengthen support for herbivory as a basal state in Cynipidae+Figitidae followed by reversal to a parasitoid lifecycle in Figitidae (Fig. 6B). On the other hand, if *Eschatocerus* were shown to be koinobiont parasitoids of other gall inhabitants, this would further strengthen the independent phytophagy scenario (Fig. 6A).

### Transitions between gall inducing and inquiline lifecycles

Whichever of our two hypotheses (gall inducers first, or inquilines first) is correct, the history of transitions between inquilinism and gall induction is clearly more complex than the origin of phytophagy. The only transition that is clearly supported by phylogenetic evidence at this point is the origin of gall induction by *Synergus itoensis* and close relatives from inquiline ancestry within the Synergini (Ide et al. 2018). If we assume that transitions have always been from inquilinism to gall induction in the Cynipidae (s. str.), then the inquilines-first scenario appears slightly more likely (Fig. 7A). However, if we assume (as seems most likely) that gall induction in the *S. itoensis* lineage represents reversal from an inquiline life cycle, then the gallers-first scenario (Fig. 7B) provides a more parsimonious explanation of the remaining transitions. Which hypothesis is better supported depends crucially on the relative ease (in evolutionary terms) or weight (in terms of inferred state changes) of transitions between the alternative states of gall induction and inquiline lifecycles (Stone & French 2003). While both gall inducers and inquiline cynipids can cause the development of nutritive gall tissues on which the larvae feed, only true inducers can cause the development of gall tissues *de novo*, and the development of the structurally complex outer gall tissues that characterize many cynipid galls. If it is easier to transition from full gall induction to a simpler inquiline life history than *vice versa*, then a gallers-first scenario may be more likely *a priori.* Alternatively, it might be a relatively minor step in evolutionary terms for cynipids to transition from inquilinism to becoming a gall inducer. We currently know too little about the differences between these alternative life histories to provide any clear weighting of transition probabilities between them, beyond suspecting that unweighted parsimony may be an unreliable guide. While the evolution of gall induction in *Synergus itoensis* shows that gall induction can evolve from inquilinism, the galls they induce consist only of nutritive tissues and lack morphologically complex non-nutritive tissues. Some Synergini inquilines do modify the complex gall morphology of host galls usurped at a very early stage in their development (Pénzes et al. 2009), but no case is yet known of a shift from inquilinism to gall induction that also includes ability to induce complex gall phenotypes.

Again, the life history of some key taxa is important in weighing these alternative scenarios. Demonstration that Eschatocerini are inquilines would strengthen the inquilines-first scenario, while demonstration that they are true gall inducers would strengthen support for the gallers-first scenario. The Qwaqwaiini is another taxon for which more detailed life-history information would be valuable. According to the only existing report it is a gall inducer (Liljeblad et al. 2011), but it remains possible that it could be an inquiline, like most members of the Synergini (s. str.) + Rhoophilini, of which we infer it to be the sister group. Such a demonstration would strengthen support for the inquilines-first scenario by removing one of the independent origins of gall inducers. Finally, we note that an ancestral state for inquilinism in Cynipidae (s. str.) also requires that the ancestral host was not itself a cynipid gall inducer. While rare examples of inquiline cynipids developing in non-cynipid galls are known (Askew 1999; van Noort et al. 2007), it is notable that the vast majority of inquiline cynipids develop in cynipid galls.

Transitions between strikingly different life histories, such as those between koinobiont endoparasitoids, gall inducers and inquilines in cynipoids, should have major effects on genomes. For instance, transitions to or from a koinobiont endoparasitic life history should involve recruitment or loss of a swathe of genes or gene functions associated, for example, with suppressing or evading host immune systems, maintaining basic physiological functions within a host body, and adjusting larval development and feeding patterns so that the host larva survives and develops normally as long as possible. This should be noticeable as an unusual number of protein-coding genes with markedly increased or decreased rates of non-synonymous rates of evolution along branches of the phylogeny involving life history changes. Similarly, the genes undergoing unusual amounts of change should also belong to particular functional categories. Transitions between gall inducers and inquilines may be less dramatic but should nevertheless leave similar genomic signatures. A recent study suggests that this is indeed the case for the transition from inquilines to gall inducers in species related to *Synergus itoensis* (Gobbo et al. 2020). Whether such genomic signatures of life history transitions can be detected deeper down in the cynipoid tree remains unclear. However, this is clearly a possibility that is well worth investigating, and the genomic data reported here represents a first step in supporting such a line of research.

## Acknowledgements

We would like to thank Jean-Yves Rasplus and Astrid Cruaud for their willingness to share barcoding and UCE data providing critical clues to the life history of *Cecinothofagus* prior to publication. Jean-Yves Rasplus also provided valuable comments on the manuscript.

## Author contributions

G.N.S. and J.H. conceived the study. J.H., J.A.N., G.K. and E.G. sequenced genomes. J.H. and E.G. assembled genomes and generated gene alignments. E.G., F.R., N.L. and A.B. designed and performed phylogenetic analyses. J.L.N.A. contributed material, life-history information and taxonomic discussion. E.G., G.N.S., J.H. and F.R. wrote the first draft of the manuscript. The final manuscript was a joint effort.

## Data availability

Raw sequencing data is available under EBI Bioprojects PRJEB13424, PRJEB45812, PRJEB51101 and PRJEB37996. Scripts, datasets, and result files are available from https://github.com/ronquistlab/cynipoid_phylogenomics.

