## Supplementary Material for "Phylogenomic Analysis of Protein-Coding Genes Resolves Complex Gall Wasp Relationships"

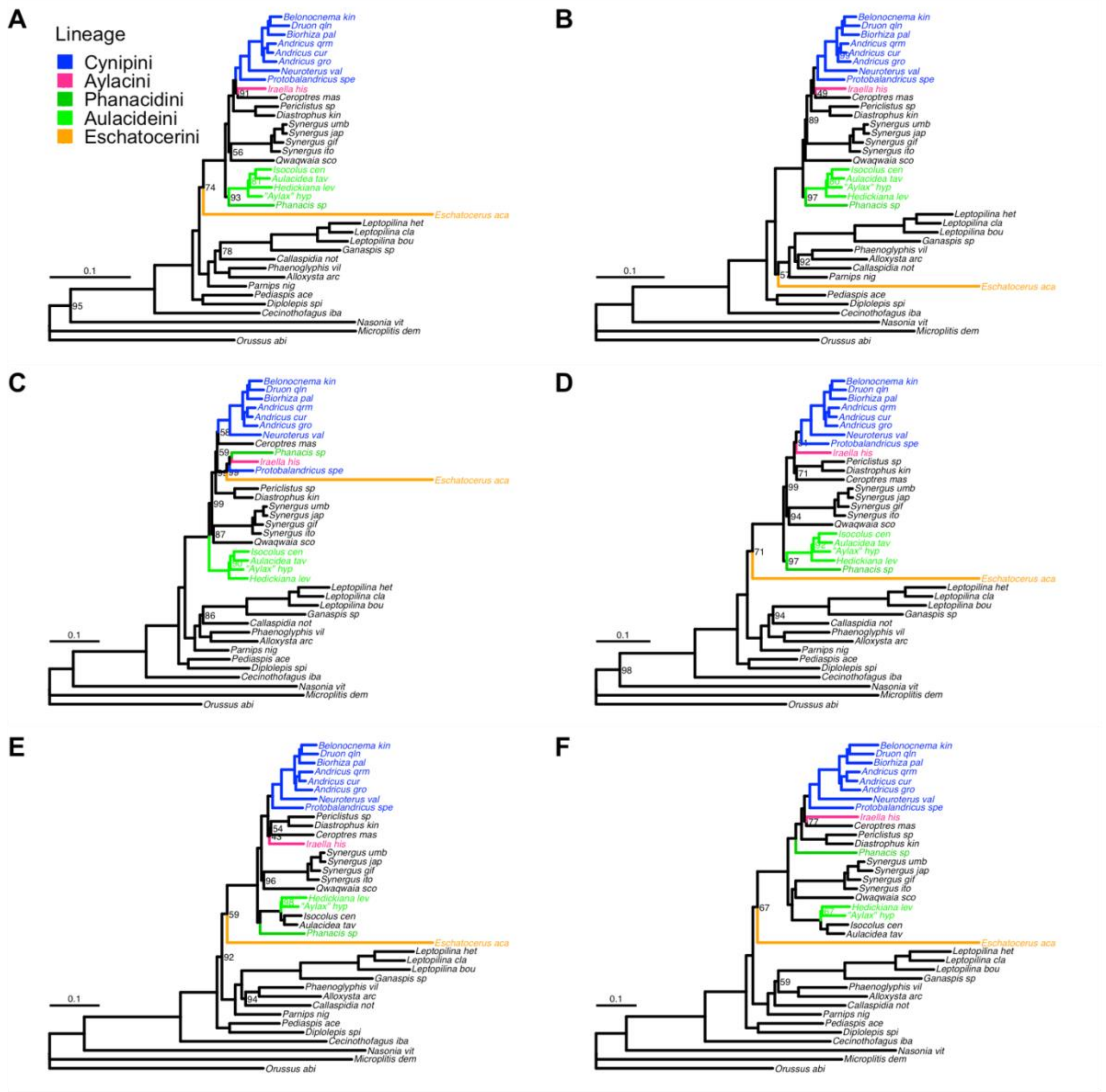


**Figure S1.** Phylogenetic results for six equally-sized, quality-ranked subsets of the T34-G542 dataset, analyzed using IQTree under the C60+I+G5 model. The raw alignments were subjected to filtering and quality ranking by HmmCleaner only. Support values (ultrafast bootstrap) are shown on branches only if they are less than 100%. **A**. Less than 1.8% of sites filtered out (best quality). **B**. From 1.8% to 3.5% filtered out. **C**. From 3.5% to 4.9% filtered out. **D**. From 4.9% to 7.4% filtered out. **E**. From 7.4% to 11% filtered out. **F**. More than 11% filtered out (worst quality). The two best subsets (A-B) yield congruent results except for the position of *Eschatocerus*, which varies slightly but without strong conflict in support values. The results for these subsets are also consistent with those from the best alignments using other filtering and quality ranking criteria (Figs. 3, S2, S3).

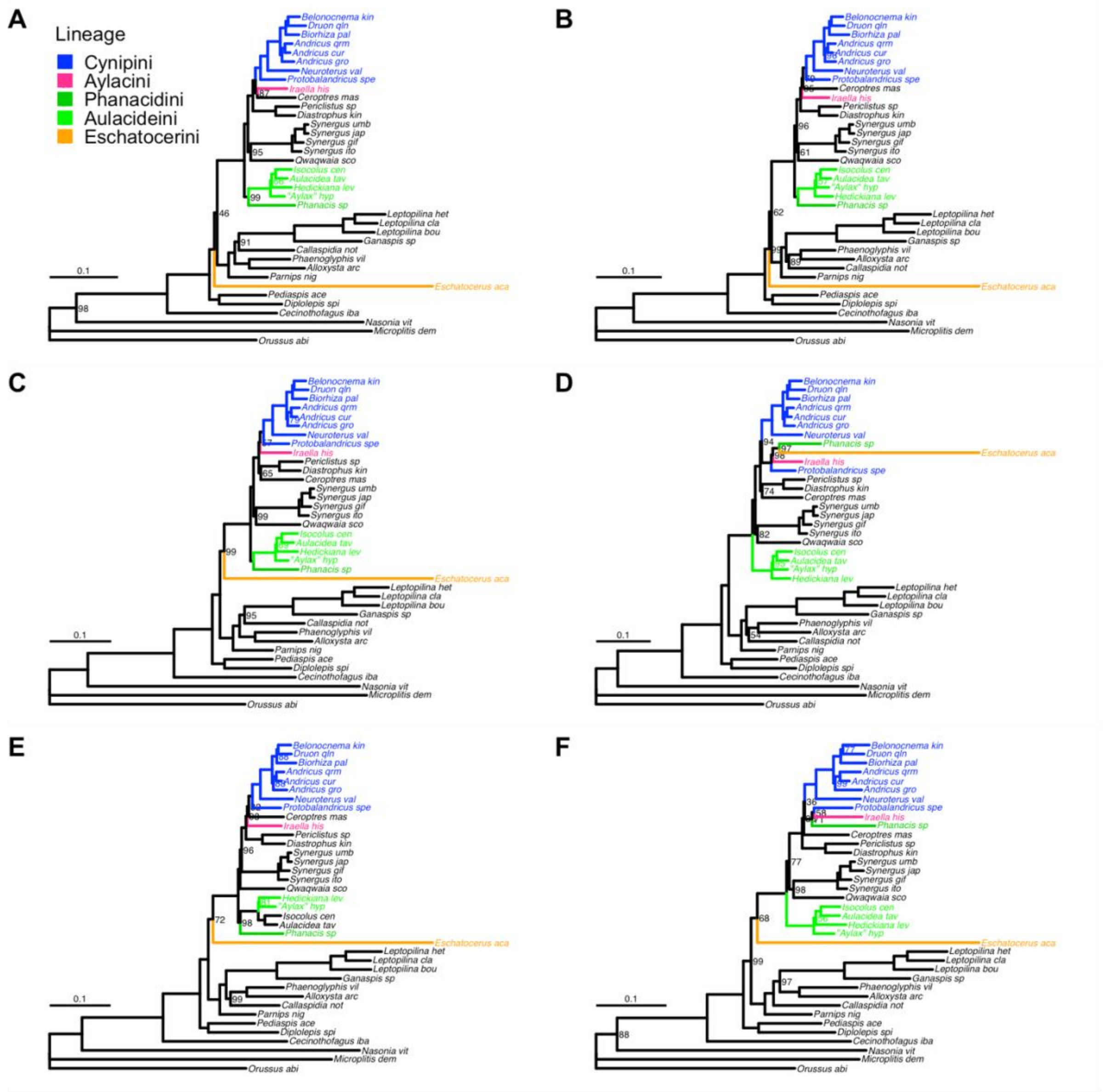


**Figure S2.** Phylogenetic results for six equally-sized, quality-ranked subsets of the T34-G542 dataset, analyzed using IQTree under the C60+I+G5 model. The raw alignments were subjected to filtering and quality ranking by GBLOCKS and HmmCleaner. Support values (ultrafast bootstrap) are shown on branches only if they are less than 100%. **A**. Less than 16.5% of sites filtered out (best quality). **B**. From 16.5% to 28.8% filtered out. **C**. From 28.8% to 39.6% filtered out. **D**. From 39.6% to 52.2% filtered out. **E**. From 52.2% to 64.1% filtered out. **F**. More than 64.1% filtered out (worst quality). The two best subsets (A-B) yield congruent results except for the position of *Eschatocerus*, which varies slightly but without strong conflict in support values. The results for these subsets are also consistent with those from the best alignments using other filtering and quality ranking criteria (Figs. 3, S1, S3).

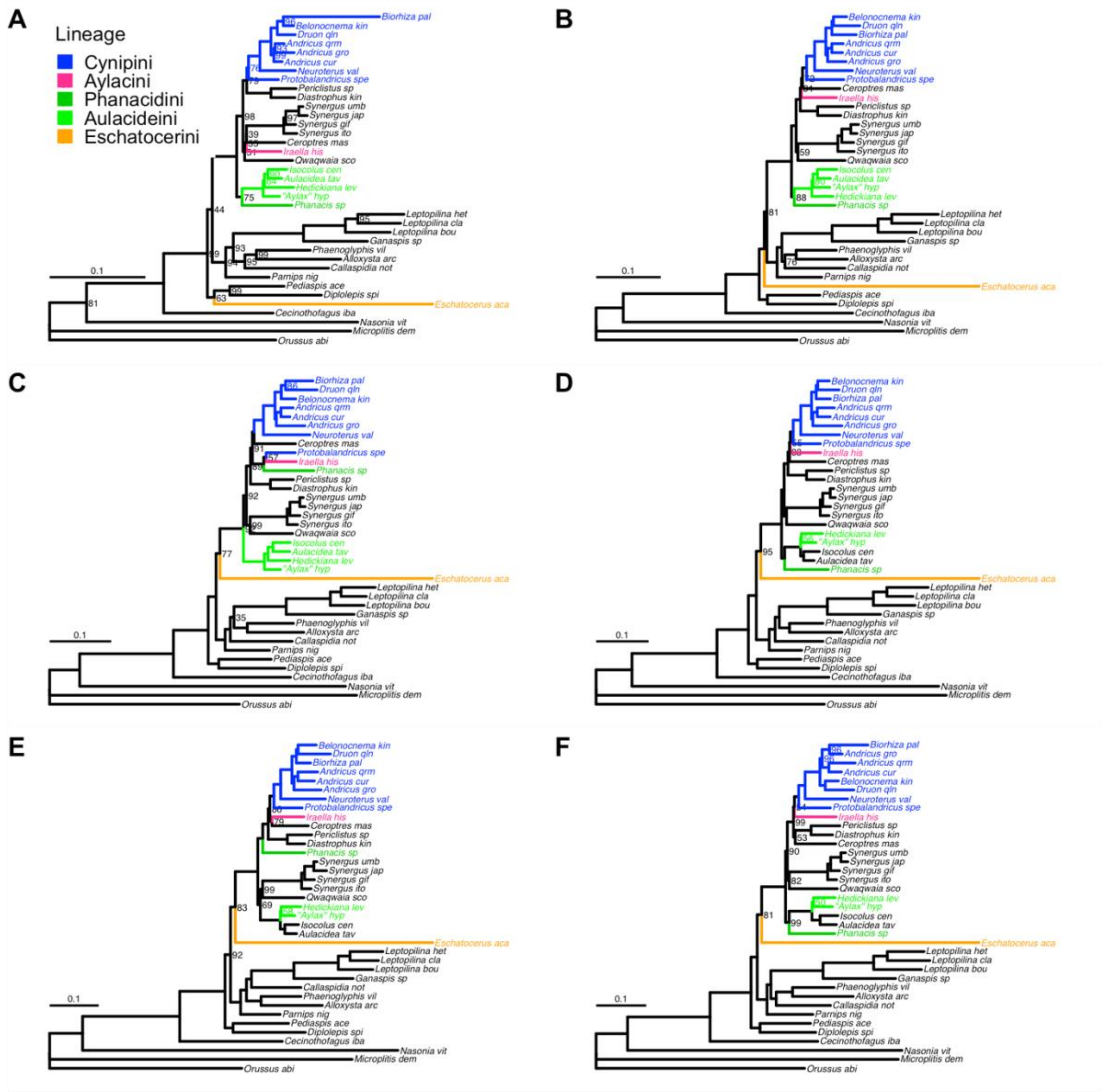


**Figure S3.** Phylogenetic results for six quality-ranked subsets of the T34-G542 dataset, analyzed using IQTree under the C60+I+G5 model. The raw alignments were subjected to filtering and quality ranking by OdSeq only. Support values (ultrafast bootstrap) are shown on branches only if they are less than 100%. **A**. No outlier sequence removed (best quality). **B**. One to three sequences removed. **C**. Four or five sequences removed. **D**. Six or seven sequences removed. **E**. Eight or nine sequences removed. **F**. Ten or more sequences removed (worst quality). Because of the discrete nature of the OdSeq filtering (number of sequences removed), the best-quality subset (A) is significantly smaller than the others, explaining the lower support values. Except for some variations with weak support, the two best subsets yield phylogenetic results that are congruent with each other and with those from the best alignments using other filtering and quality ranking criteria (Figs. 3, S1, S2).

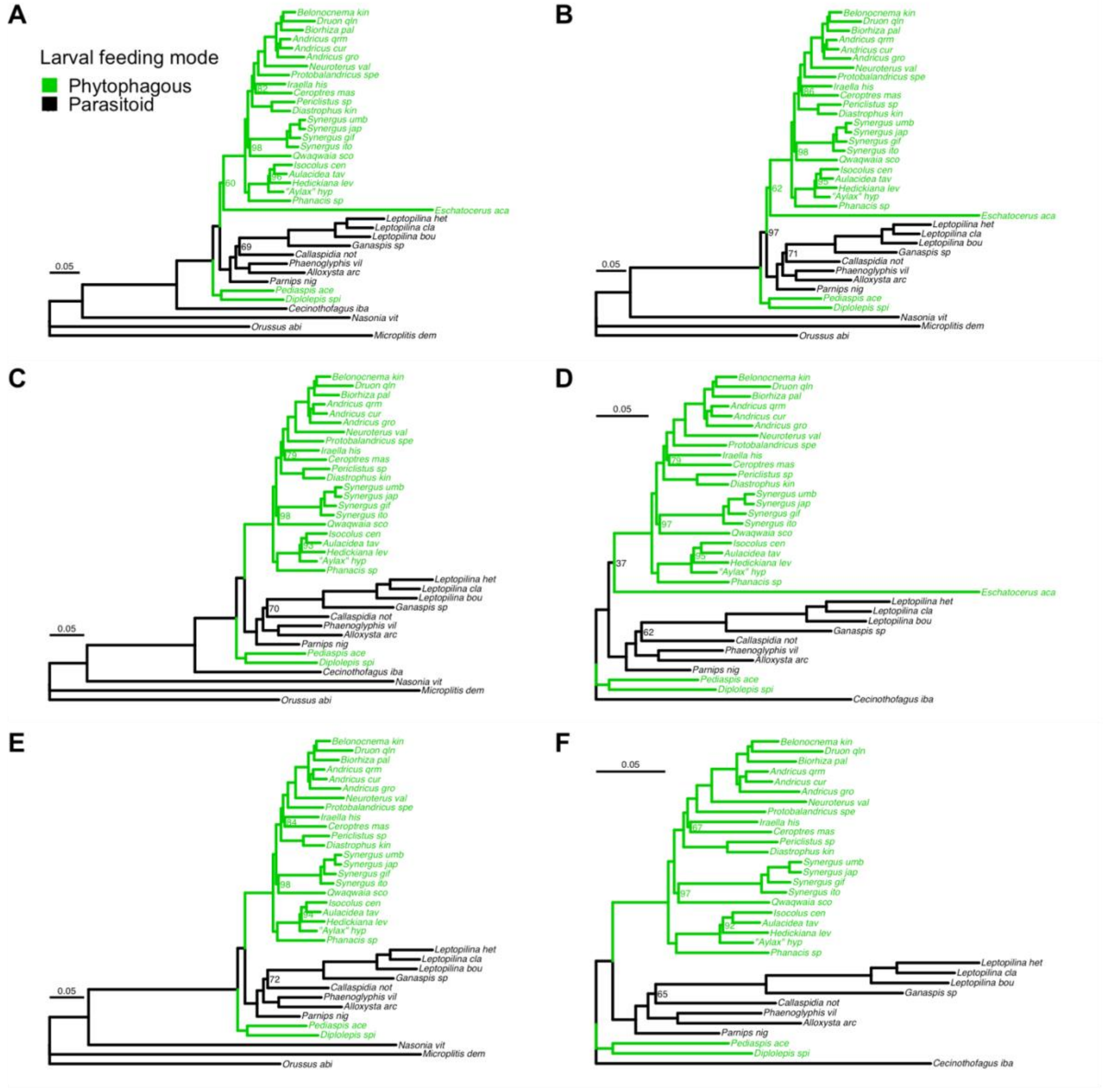


**Figure S4**. Testing the potential effect of long-branch taxa on phylogenetic results using IQTree under the C60+I+G5 model rather than PhyloBayes. Otherwise, this is an identical replicate of the analyses shown in the Main Text (Fig. 4). Branch support values (ultrafast bootstrap) are only shown if they are less than 100%. **A**. Analysis of the full taxon set. **B**. *Cecinothofagus* excluded. **C**. *Eschatocerus* excluded. **D**. Outgroups excluded. **E**. *Cecinothofagus* and *Eschatocerus* excluded. **F**. *Eschatocerus* and outgroups excluded. Results are identical to those obtained with PhyloBayes and the CAT-F81 model (Fig. 4), except a minor and weakly supported difference in the positions of *Ceroptres* and *Iraella* when *Eschatocerus* and outgroups are excluded (F).

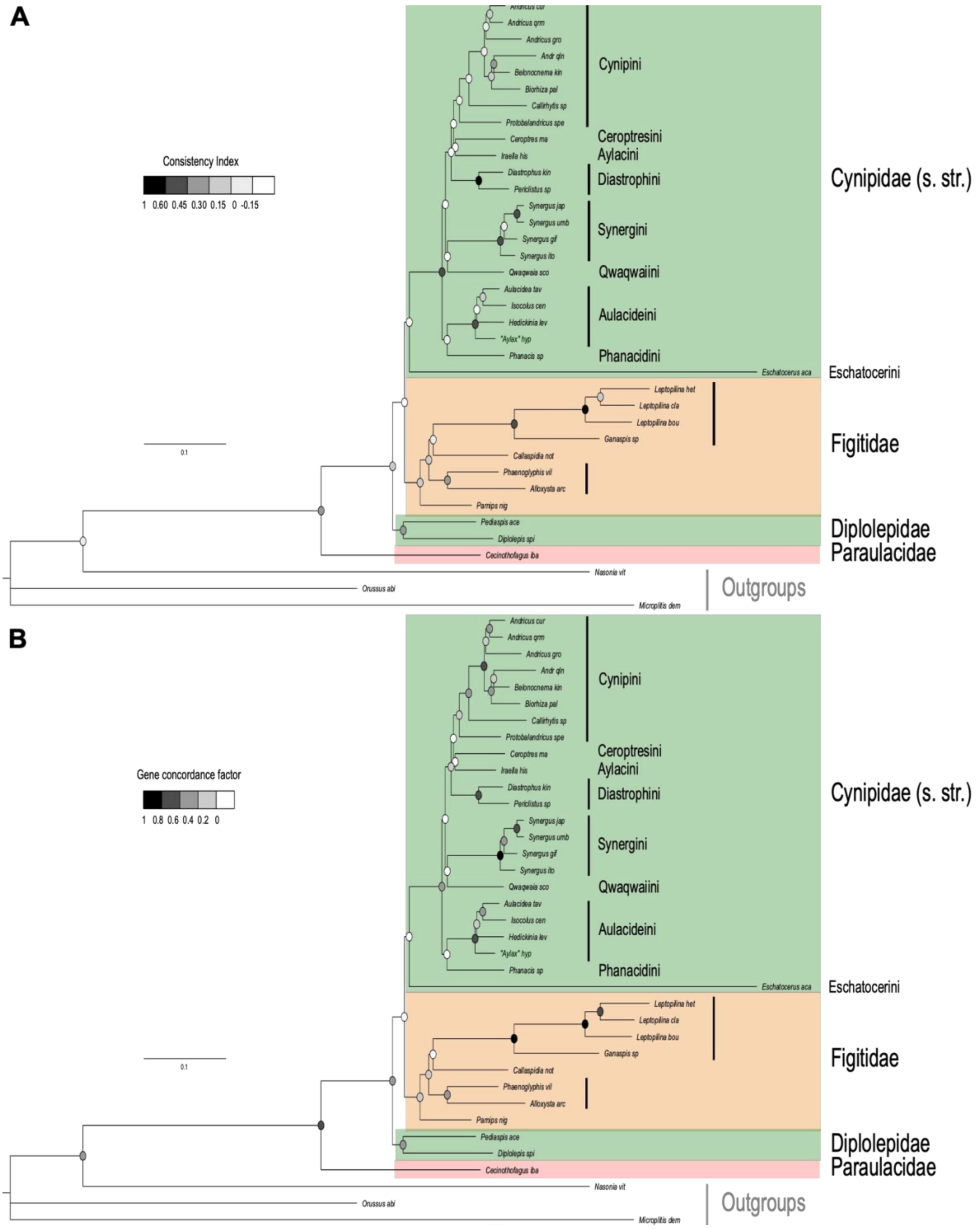


**Figure S5**. Results from gene tree concordance analysis. **A**. Gene tree consistency indices. **B**. Gene tree concordance factors.

**Table S1. Species metadata and assembly statistics for the genomes and transcriptomes used in this study**. Columns contain information on collection location where available, whether the data was assembled here or previously, and links to raw read data. Genome/transcriptome assembly quality metrics include complete and partial BUSCO (v4.0.6) scores for both Hymenoptera and Eukaryota datasets respectively, assembly size, N50 and number of contigs, repeated for contigs >200 bp.

[Table provided in a separate Excel file]

**Table S2**. **Completeness of different BUSCO gene sets per taxon**. The table gives the number of genes of each gene set present in each taxon, as well as the proportion of the total set being present in the taxon. Taxon names correspond to those used in the datasets. Note that, because of taxonomic changes or incorrect annotations in public databases, Beloc_tr is used for *Belonocnema kinseyi*, Calli_sp for *Neuroterus valhalla* and Andr_qln for *Druon quercuslanigerum*.

| | **Gene set (minimum number of taxa)** | | | | | | | | |
|---|---|---|---|---|---|---|---|---|---|
| **Taxon** | **37** | **36+** | **35+** | **34+** | **33+** | **32+** | **31+** | **30+** | **All** |
| Allox_ar | 31 (100%) | 123 (100%) | 293 (99%) | 536 (99%) | 797 (98%) | 1076 (97%) | 1432 (96%) | 1807 (95%) | 4730 (80%) |
| Andr_cur | 31 (100%) | 123 (100%) | 295 (100%) | 538 (99%) | 807 (99%) | 1098 (98%) | 1460 (98%) | 1838 (97%) | 4729 (80%) |
| Andr_gro | 31 (100%) | 123 (100%) | 290 (98%) | 526 (97%) | 763 (94%) | 992 (89%) | 1254 (84%) | 1506 (79%) | 2872 (49%) |
| Andr_qln | 31 (100%) | 122 (99%) | 294 (99%) | 530 (98%) | 778 (95%) | 1037 (93%) | 1324 (89%) | 1628 (86%) | 3214 (55%) |
| Andr_qrm | 31 (100%) | 119 (97%) | 280 (95%) | 500 (92%) | 750 (92%) | 1002 (90%) | 1318 (88%) | 1645 (87%) | 4015 (68%) |
| Aula_tav | 31 (100%) | 123 (100%) | 295 (100%) | 538 (99%) | 802 (98%) | 1092 (98%) | 1457 (97%) | 1837 (97%) | 4515 (77%) |
| Aylax_hy | 31 (100%) | 123 (100%) | 296 (100%) | 542 (100%) | 813 (100%) | 1112 (100%) | 1489 (100%) | 1878 (99%) | 4947 (84%) |
| Beloc_tr | 31 (100%) | 122 (99%) | 293 (99%) | 536 (99%) | 800 (98%) | 1090 (98%) | 1454 (97%) | 1843 (97%) | 5388 (91%) |
| Bior_pal | 31 (100%) | 85 (69%) | 158 (53%) | 268 (49%) | 398 (49%) | 508 (46%) | 660 (44%) | 797 (42%) | 1931 (33%) |
| Callas_no | 31 (100%) | 121 (98%) | 281 (95%) | 490 (90%) | 691 (85%) | 882 (79%) | 1092 (73%) | 1281 (67%) | 2076 (35%) |
| Calli_sp | 31 (100%) | 123 (100%) | 293 (99%) | 528 (97%) | 777 (95%) | 1053 (94%) | 1369 (92%) | 1682 (89%) | 3366 (57%) |
| Cecin_ib | 31 (100%) | 117 (95%) | 263 (89%) | 431 (80%) | 574 (70%) | 686 (62%) | 805 (54%) | 910 (48%) | 1376 (23%) |
| Cerop_ma | 31 (100%) | 123 (100%) | 293 (99%) | 532 (98%) | 795 (98%) | 1074 (96%) | 1424 (95%) | 1796 (95%) | 3982 (68%) |
| Diast_ki | 31 (100%) | 123 (100%) | 295 (100%) | 532 (98%) | 785 (96%) | 1062 (95%) | 1399 (94%) | 1741 (92%) | 3815 (65%) |
| Dipl_spi | 31 (100%) | 123 (100%) | 292 (99%) | 532 (98%) | 798 (98%) | 1085 (97%) | 1433 (96%) | 1785 (94%) | 4041 (69%) |
| Escha_ac | 31 (100%) | 123 (100%) | 291 (98%) | 514 (95%) | 755 (93%) | 1021 (92%) | 1350 (90%) | 1682 (89%) | 3974 (67%) |
| Ganas_sp | 31 (100%) | 106 (86%) | 238 (80%) | 386 (71%) | 584 (72%) | 788 (71%) | 1046 (70%) | 1322 (70%) | 3431 (58%) |
| Hedic_le | 31 (100%) | 123 (100%) | 295 (100%) | 536 (99%) | 799 (98%) | 1089 (98%) | 1444 (97%) | 1808 (95%) | 4106 (70%) |

| | | | | | | | | | |
|---|---|---|---|---|---|---|---|---|---|
| Irae_his | 31 (100%) | 118 (96%) | 247 (83%) | 405 (75%) | 529 (65%) | 633 (57%) | 737 (49%) | 827 (44%) | 1090 (19%) |
| Isoc_cen | 31 (100%) | 123 (100%) | 295 (100%) | 534 (99%) | 798 (98%) | 1084 (97%) | 1441 (96%) | 1812 (95%) | 4163 (71%) |
| Lepto_bo | 31 (100%) | 121 (98%) | 291 (98%) | 527 (97%) | 787 (97%) | 1067 (96%) | 1431 (96%) | 1812 (95%) | 5110 (87%) |
| Lepto_cl | 31 (100%) | 123 (100%) | 293 (99%) | 533 (98%) | 795 (98%) | 1088 (98%) | 1448 (97%) | 1833 (97%) | 4726 (80%) |
| Lepto_he | 31 (100%) | 123 (100%) | 291 (98%) | 537 (99%) | 801 (98%) | 1090 (98%) | 1455 (97%) | 1831 (96%) | 5028 (85%) |
| Parn_nig | 31 (100%) | 122 (99%) | 295 (100%) | 532 (98%) | 802 (98%) | 1093 (98%) | 1461 (98%) | 1848 (97%) | 4735 (80%) |
| Pedia_ac | 31 (100%) | 123 (100%) | 291 (98%) | 526 (97%) | 772 (95%) | 1029 (92%) | 1351 (90%) | 1670 (88%) | 3572 (61%) |
| Peric_sp | 31 (100%) | 121 (98%) | 291 (98%) | 530 (98%) | 794 (97%) | 1082 (97%) | 1438 (96%) | 1794 (94%) | 4512 (77%) |
| Phaen_vi | 31 (100%) | 121 (98%) | 287 (97%) | 520 (96%) | 769 (94%) | 1041 (93%) | 1382 (92%) | 1721 (91%) | 3807 (65%) |
| Phana_sp | 31 (100%) | 121 (98%) | 271 (92%) | 470 (87%) | 661 (81%) | 842 (76%) | 1007 (67%) | 1172 (62%) | 1894 (32%) |
| Prot_spe | 31 (100%) | 117 (95%) | 282 (95%) | 505 (93%) | 729 (89%) | 969 (87%) | 1230 (82%) | 1483 (78%) | 2555 (43%) |
| Qwaq_sco | 31 (100%) | 123 (100%) | 294 (99%) | 533 (98%) | 793 (97%) | 1074 (96%) | 1428 (96%) | 1790 (94%) | 4143 (70%) |
| Syne_gif | 31 (100%) | 123 (100%) | 293 (99%) | 534 (99%) | 798 (98%) | 1094 (98%) | 1467 (98%) | 1858 (98%) | 5303 (90%) |
| Syne_ito | 31 (100%) | 123 (100%) | 294 (99%) | 535 (99%) | 796 (98%) | 1089 (98%) | 1455 (97%) | 1849 (97%) | 5161 (88%) |
| Syne_jap | 31 (100%) | 123 (100%) | 294 (99%) | 534 (99%) | 800 (98%) | 1093 (98%) | 1462 (98%) | 1851 (97%) | 5322 (90%) |
| Syne_umb | 31 (100%) | 122 (99%) | 293 (99%) | 538 (99%) | 810 (99%) | 1104 (99%) | 1477 (99%) | 1873 (99%) | 5354 (91%) |
| Nasonia | 31 (100%) | 122 (99%) | 291 (98%) | 525 (97%) | 789 (97%) | 1080 (97%) | 1452 (97%) | 1844 (97%) | 5675 (96%) |
| Orussus | 31 (100%) | 122 (99%) | 293 (99%) | 535 (99%) | 802 (98%) | 1096 (98%) | 1471 (98%) | 1869 (98%) | 5743 (98%) |
| Micropl | 31 (100%) | 123 (100%) | 293 (99%) | 530 (98%) | 796 (98%) | 1092 (98%) | 1464 (98%) | 1864 (98%) | 5732 (97%) |
| **Total** | **31 (100%)** | **123 (100%)** | **296 (100%)** | **542 (100%)** | **815 (100%)** | **1115 (100%)** | **1495 (100%)** | **1899 (100%)** | **5890 (100%)** |
